# Neural mechanisms by which spaced learning promotes understanding

**DOI:** 10.1101/2021.09.06.459209

**Authors:** Yaowen Liang, Yuying He, Zijian Cheng, Yiwen Li, Linyan Liu, Yanhong Wu, Yufeng Sun, Fujing Ren, Meiqiong Shuai, Wenbo Luo, Xiujie Yang, Liang Zhang

## Abstract

Spaced learning reliably improves memory. A few studies suggest it also benefits understanding and problem solving, but none has examined the neural mechanism. We hypothesised that this benefit arises because spacing changes the memory trace rather than merely strengthening it. To test this, we taught participants how unfamiliar words’ meanings could be derived from familiar parts: meaningful rather than rote learning. Each participant learned words in one session, or in two sessions about 2 h (massed) or 48 h (spaced) apart, to the same criterion; one-session words served as the reference. A common interval separated the last session from a delayed test with electroencephalography (EEG). Immediately after learning, self-rated understanding did not differ between schedules, yet under spacing participants revised more of their written test responses between sessions (35% versus 25%). At the delayed test, the spacing effect appeared: recognition was higher and self-rated explanatory retrieval more fluent after spaced than after massed learning. At the neural level, late-retrieval EEG patterns for remembered words were less similar to the reference after spaced than after massed learning, over right frontocentral and temporal sensors. In partly overlapping sensors and windows, lower similarity was statistically associated with a larger fluency advantage, suggesting that the change is directed rather than random. Behavioural evidence hinted at convergence across learners: between sessions, revisions under spacing were smaller and moved responses toward those of others. Together, these findings suggest that spacing changes the memory trace rather than merely strengthening it, in a direction linked to self-rated understanding.

**Significance Statement:** Spacing out study helps memory, but any benefit for understanding may not show when study ends. Each learner studied how familiar parts explained unfamiliar words’ meanings, revisiting some after about two hours and others after about two days. Both schedules ended at the same mastery criterion, with similar ratings of understanding. Yet with the longer gap, learners had revised written explanations more often, in smaller steps toward other learners’ wording. After an equal delay, they rated those explanations easier to reconstruct. Brain activity during retrieval was consistent with a memory that spacing had reorganised, not merely strengthened, in a way linked to those ratings. The moment a learner feels they understand may be too early to judge a study schedule.

## Introduction

Spacing is one of the most reliable ways to improve delayed retention (Cepeda et al., 2006; Latimier et al., 2021). Learning, however, demands more than retention: its value lies in building the relations and explanations that make knowledge meaningful and generative (Mayer, 2002; Spens & Burgess, 2024). Yet laboratory research on spacing has long focused primarily on retaining learned material (Cepeda et al., 2006; Gerbier & Toppino, 2015). To maximise experimental control, these studies have often deliberately avoided meaningfully structured materials while tightly controlling the timing and duration of learning. This emphasis on control is particularly pronounced in neural studies. Such tight control over both the materials and the learning process risks obscuring how spacing operates in meaningful learning. By contrast, educational research has begun to extend the study of spacing beyond rote retention to meaningful learning (Vlach & Sandhofer, 2012; see also Samani & Pan, 2021, for problem solving under interleaved practice). However, the neural basis of these benefits has not been investigated. To our knowledge, the present study is the first to examine whether spaced restudy improves explanatory understanding compared with massed restudy and how any such improvement is reflected in neural representations.

To explain the potential benefits of spacing for understanding, we first consider theories of how spacing improves memory. A central question is whether repeated learning reinforces an existing memory representation or changes its organisation. Greater neural pattern similarity across repeated presentations has been associated with better subsequent memory in both functional MRI (fMRI) and electroencephalography (EEG) studies (Lu et al., 2015; Xue et al., 2010). Feng et al. (2019) extended this finding to spacing, linking its memory benefit to greater pattern similarity and interpreting this as reinstatement and strengthening of prior representations. Other theories, by contrast, hold that repeated learning changes the memory rather than merely strengthening it (Glenberg, 1979; Lee, 2009; Moscovitch & Gilboa, 2024; Smith & Scarf, 2017). For meaningful learning, we favour this second view: understanding concerns how elements of knowledge are related, not simply how readily they can be recalled. Such change could itself take two forms: encoding variability theory emphasises the accumulation of varied contextual details across learning encounters, providing additional routes to retrieval (Glenberg, 1979; Maddox, 2016; Melton, 1970), whereas transformation and reorganisation accounts emphasise restructuring the relations among the elements of a memory. Rather than adjudicate between them, we ask whether the change has a direction: whether it moves the memory toward meaning and tracks explanatory understanding. We outline below how our study examines this question.

Doing so first requires materials in which understanding can be assessed through the reconstruction of an explanatory relation. Much of the spacing literature has used materials such as word pairs and isolated facts, for which successful recall does not require reconstructing an internal explanatory relation (Carpenter et al., 2022; Cepeda et al., 2006). Some spacing research has added externally supplied semantic contexts (Dempster, 1987), while related vocabulary-learning work has used mnemonic keywords (McDaniel & Tillman, 1987); neither requires learners to reconstruct relations intrinsic to the material. By contrast, etymology-based vocabulary learning asks learners to connect a word’s form and etymological cues to its meaning, so that the relation to be reconstructed is intrinsic to the material (Hosseini et al., 2012; Templeton, 2012), in line with the classical sense of meaningful learning (Mayer, 2002). The task therefore distinguishes knowing a word’s translation from recovering the relation that makes it intelligible, allowing us to test whether spacing affects how fluently the relation can be reconstructed, rather than only whether the answer is retained.

Doing so also requires a neural measure for which reinforcement and change make different predictions. The typical approach has been to compare neural patterns between the first and second encoding episodes (E1–E2 similarity; Feng et al., 2019; Zou et al., 2025). This approach has the merit of observing the repeated encounter directly. However, on the study-phase retrieval account (Benjamin & Tullis, 2010; Maddox et al., 2018), the second episode necessarily involves retrieving the first, so encoding and retrieval are intermixed within it; as a result, the same E1–E2 similarity can support opposite interpretations. If spacing produces greater similarity across encoding episodes, this could reflect successful retrieval of the prior trace, or forgetting-induced re-encoding under similar task constraints—because knowledge structures are relatively stable (cf. Hanfstingl et al., 2021), participants are likely to recruit the same knowledge and processes for the same task, producing similar patterns even without retrieving the prior trace. Because the relative contributions of retrieval and re-encoding cannot be directly measured, E1–E2 similarity cannot cleanly adjudicate between these two readings. A small body of work has addressed this by shifting the neural observation window to the retrieval stage (Li & Yang, 2020; Takashima et al., 2007; Yang et al., 2025). But Takashima et al. (2007) and Li & Yang (2020) contrasted massed with spaced learning directly, yielding a difference without a direction. Yang et al. (2025) went a step further, reading later retrievals against an earlier one across three sessions (immediate, 1-week, and 1-month delays), which restores a directional reference; but because the same items were retrieved at each session, and each retrieval is a learning event—the testing effect (Roediger & Karpicke, 2006)—that reference is contaminated, conflating the original spacing manipulation with the cumulative effects of prior retrieval. What none provides is a clean anchor—material studied once and not subjected to an earlier delayed test— against which spaced and massed restudy can each be read, without contamination, as reorganising the original trace or leaving it intact.

The present study was designed to provide such an anchor. We added a control condition (O): a separate set of words, independent of the spaced and massed sets, that received only one study episode without any subsequent review. This set provided a once-learning reference against which the spaced and massed retrieval patterns could be read as departures — a reference that makes reorganisation of the trace visible, not a neutral zero. Each item, including those in the reference set, contributed to only one delayed EEG retrieval assessment, avoiding cumulative retrieval-practice effects across repeated delayed tests. The sole once-learning episode and the second learning episodes for both repeated conditions all occurred on Day 3, so all three conditions had the same interval between their last study episode and the Day 5 test, at which neural pattern similarity (NPS) was measured. By recording EEG during the final retrieval attempt, when all three conditions faced the same reconstruction demand, we could read NPS as reflecting retrieval after different learning histories, rather than a mixture of encoding and retrieval that differs between conditions. The same design also removed the constraint on learning time noted at the outset. E1–E2 comparisons require matched observation windows, which limits how long each learning encounter can run. Treating those matched windows as equal amounts of learning is natural under incremental-strengthening models (cf. Pavlik & Anderson, 2005), but equal time need not mean that the material is equally well learned: understanding depends on organising and integrating relations rather than exposure alone (Mayer, 2002). A fixed interval could stop one schedule before the relation was mastered while forcing another to continue beyond that point, where additional study can yield sharply diminishing long-term returns (Rohrer & Pashler, 2007). Clock time is therefore a poor proxy for a matched learning endpoint. Because we measured at final retrieval rather than across encoding episodes, we could end learning at the same mastery criterion instead: participants studied until they judged the relation understood well enough, with an immediate mastery test providing an objective check on that judgement. By controlling apparent mastery rather than clock time, we could ask what remains after learners appear to have understood.

Together, these controls limit how a later neural difference can be explained. The two schedules ended learning at the same mastery criterion, were tested after the same delay, and contributed only successfully remembered items. A later difference would therefore be hard to attribute solely to how well the material was first learned, how long ago it was learned, or whether it could still be retrieved. It would instead speak to how the accessible trace was organised. We expected the second learning episode to be when the two schedules began to organise the trace differently. When the first trace was re-encountered, the longer spaced interval allowed more time for consolidation and re-encoding of the trace (Smith & Scarf, 2017), whereas massed restudy could still draw on a fresher trace of the first episode (Hanfstingl et al., 2021). The common Day 3-to-Day 5 delay then provided a test of that organisation: if the two learning histories had left differently organised traces at that endpoint, final retrieval would read out what survived after the same delay. We predicted that spacing would produce greater differentiation from the once-learning reference than massing, and that this spacing-specific differentiation would be associated with the relative delayed advantage in self-rated explanatory retrieval fluency. On a reinforcement account, by contrast, spaced restudy would only stabilise the retrieval pattern that a single episode leaves, so spaced items should resemble the once-learning reference at least as closely as massed items, with closer resemblance accompanying better retrieval. Because reconstructing an explanatory relation requires semantic and episodic information to be recovered and integrated, processes that unfold in the later part of the retrieval epoch (Johnson et al., 1998b; Moccia et al., 2025), we predicted that this differentiation would be most evident late in the epoch. We therefore targeted a bilateral central domain over fronto-central, temporal and centro-parietal sensors, which overlie regions implicated in this recovery, with the anterior temporal lobe engaged early and angular gyrus involvement emerging later (Farahibozorg et al., 2022), across the full post-stimulus epoch. If this differentiation reflects a shift of the trace toward meaning, retrieval patterns should also lean less on the visual form of the word; in a separate exploratory analysis we therefore asked whether final-test patterns still corresponded to a visual word-recognition model at early stages (Hauk et al., 2006; Laszlo & Federmeier, 2014), under each schedule. A change with a direction raises one further question: whether different learners move in the same direction. The written responses at the two initial tests offered a behavioural view of this: because participants wrote out the intermediary words and the definition each time, we could ask how often a word’s response was revised between the tests, how much it changed, and whether it became more similar to what other learners wrote for the same word.

## Results

Figure 1 shows the learning schedules and the delayed final test; Figure 2 outlines the NPS pipeline, which is described in detail in Methods.

**Figure 1.**
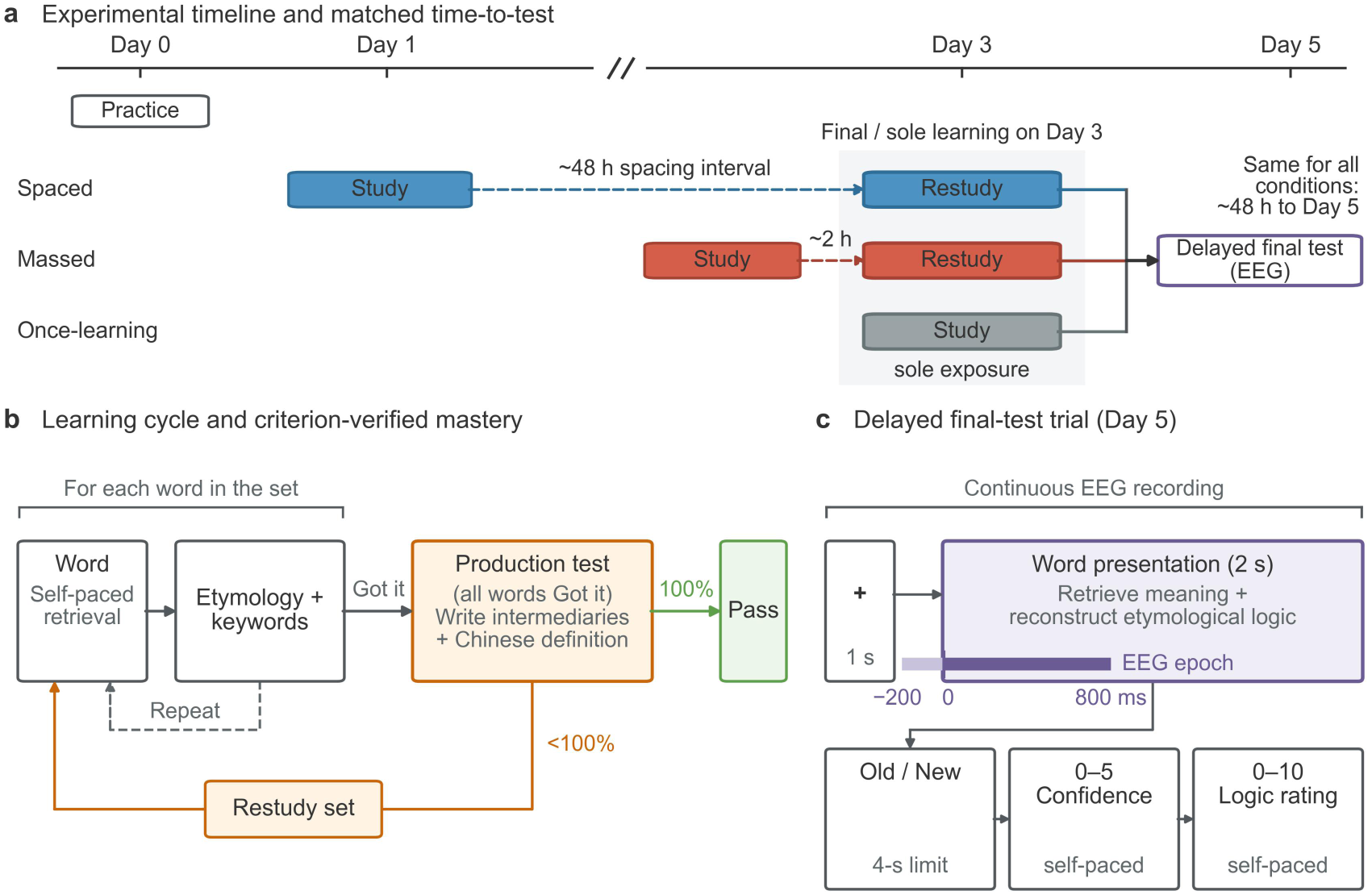
Experimental paradigm. (a) Experimental timeline. Participants practised on Day 0, then studied words under spaced (∼48 h interval), massed (∼2 h interval), and once-learning conditions, followed by a delayed final test with EEG recording on Day 5. (b) Learning phase. For each word, participants first attempted retrieval, then studied etymology and keywords, repeating until mastery before the initial test. (c) Delayed EEG final test. During the 2-s word presentation, participants attempted to retrieve the learned meaning and reconstruct the etymological logic before making an old–new recognition judgement (4-s response limit), followed by self-paced confidence and explanatory retrieval-fluency ratings.

**Figure 2.**
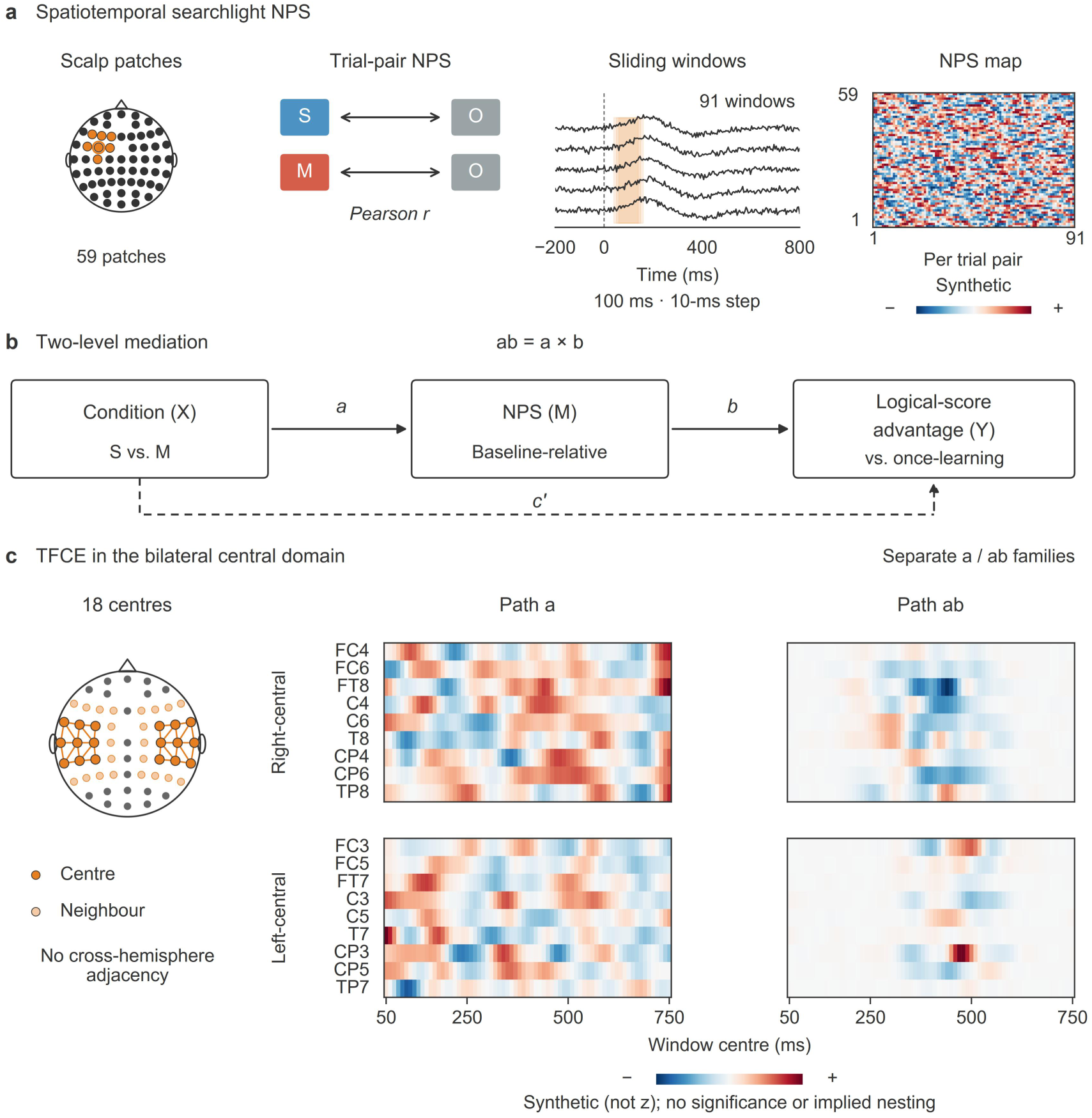
NPS analysis pipeline. (a) Spatiotemporal searchlight NPS: 59 scalp patches (each an electrode and its neighbours within 5 cm) and 91 temporal sliding windows (100 ms width, 10 ms step, −200 to 800 ms; window centres −150 to 750 ms), with pairwise Pearson r computed between each spaced (S) or massed (M) trial and once-learning (O) baseline trials; the complete 59 × 91 maps were retained for descriptive visualisation. (b) Two-level mediation model fitted by robust maximum likelihood (MLR): condition (X, spaced vs. massed) predicts the baseline-relative self-rated explanatory retrieval-fluency advantage (Y) via NPS (M), using 63,250 complete-case trial-pair rows nested within 28 participants; path a links condition to NPS, path b links NPS to the outcome, c′ is the direct effect of condition on the outcome, and the indirect effect is ab = a × b. (c) Formal inference covered the bilateral central domain of 18 searchlight centres (nine per hemisphere over the fronto-central, temporal and centro-parietal rows) at all 71 post-stimulus window centres (50–750 ms) and used threshold-free cluster enhancement (TFCE; height exponent H = 2, extent exponent E = 1) with 1,991 within-participant, intact-target-block label permutations and shared raw-maximum correction within the path-a and path-ab families.

### Behavioural Results

The behavioural results revealed a dissociation between immediate and delayed performance. At the second-session immediate test (T2), spaced and massed learning yielded comparable immediate logical scores despite longer study in the spaced condition (Figure 3a–b). These comparable scores were nonetheless accompanied by a difference in what participants wrote. Spaced learning raised the odds that a word’s first-session written response was revised at the second test (35.2% versus 24.9% of observable words; adjusted odds ratio 1.75, 95% CI [1.44, 2.13], *P* < 0.001), whereas the revisions that did occur were smaller (adjusted spaced-minus-massed difference in text-embedding distance between the two responses −0.021 [−0.037, −0.005], *P* = 0.009; Figure 3c). Revised responses also moved closer to the other participants’ formulations after spaced than after massed learning: the change from the first-session test (T1) to T2 in distance to the same-test group centre differed by −0.027 [−0.047, −0.007] (*P* = 0.007; unadjusted means: spaced −0.010, massed +0.012; Figure 3f; Supplementary Results S7). At the delayed final test, spaced learning outperformed massed learning in self-rated explanatory retrieval fluency and recognition (Figure 3d–e).

**Figure 3.**
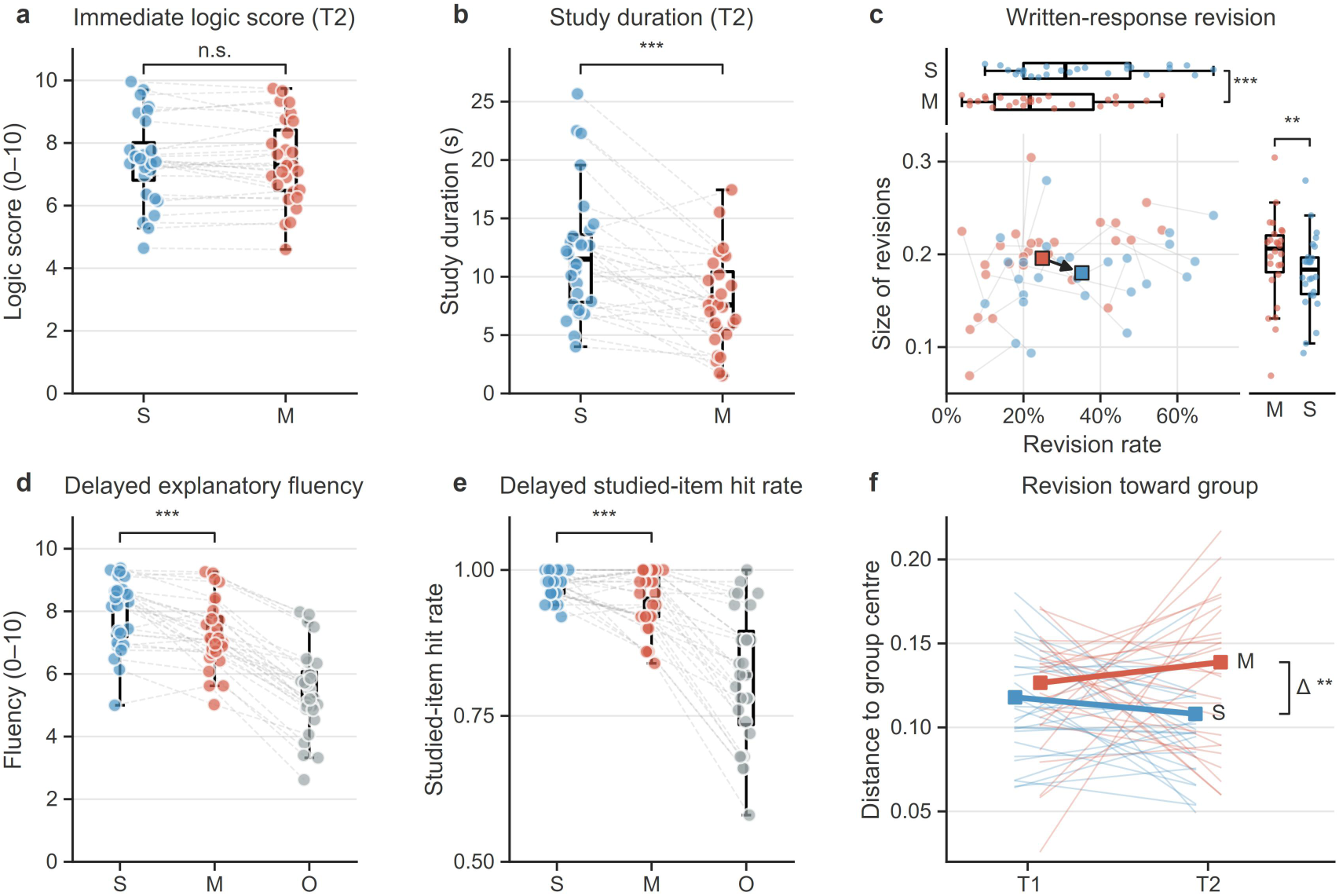
Behavioural outcomes across learning and delayed retrieval. (a) Immediate self-rated logic score at T2. (b) Study duration to mastery at T2. (c) Written-response revision between T1 and T2. The plane plots each participant’s revision rate (proportion of observable words whose written response changed) against the mean size of those revisions (text-embedding distance between the T1 and T2 responses; Methods), with the condition means joined by an arrow from massed to spaced; the margins show the rate (top) and size (right) distributions. (d) Delayed self-rated explanatory retrieval fluency. (e) Delayed studied-item hit rate. (f) Revision toward the group. For revised words, the distance between a participant’s response and the centre of the other participants’ responses to the same word at the same test (T1 against the T1 centre, T2 against the T2 centre); thin lines are participant means, bold lines word-level group means. Inner boxes show medians and interquartile ranges, coloured dots show participant summaries, and thin grey lines connect within-participant observations (spaced = blue, massed = red, once-learning = grey). Panels a, b, d and e include 28 participants; panels c and f include 26, because two participants had fewer than 20 words with both responses observed in at least one condition, an outcome-blind coverage rule. Brackets mark the primary spaced-versus-massed comparison in each panel: a–b, paired tests (one contrast at T2); c, word-level mixed models with participant and word random intercepts and covariates (rate, logistic; size, linear); d–e, covariate-adjusted trial-level models (d, linear mixed model; e, generalised estimating equations, GEE); f, the same word-level mixed model on the T1-to-T2 change (Δ). *** P < 0.001, ** P < 0.01, n.s. not significant; estimates and confidence intervals are given in the text. Both repeated schedules exceeded once-learning in d and e (Supplementary Results S4). Raw points are unadjusted participant summaries. Revision in c and f denotes observable written-response change and does not establish semantic improvement.

#### Comparable Initial Logical Scores Precede Later Explanatory Retrieval Differences

Under the criterion-based stopping rule, all condition-specific study sets reached 100% immediate production accuracy. At the first-session test (T1), logical scores showed no significant omnibus differences across conditions (*χ*²(2) = 2.62, *P* = 0.270; Supplementary Table 1), whereas study durations differed significantly (*χ*²(2) = 27.64, *P* < 0.001; Supplementary Table 1), consistent with a practice effect across the learning sequence. At T2 (only S and M conditions; O had only one scheduled learning episode), spaced learning required longer study duration than massed learning (*t*(27) = 5.07, *P* < 0.001, *d*_z_ = 0.96; Figure 3b), but did not show higher immediate logical scores (second-session logical score: *t*(27) = 0.68, *P* = 0.504, *d*_z_ = 0.13; Figure 3a).

At final test, the covariate-controlled trial-level models provided the primary test of the spacing advantage on final logical score, the self-rated explanatory retrieval fluency measure (Figure 3d), and on recognition. Controlling first-session study duration and first-session logical score and evaluating condition effects at mean College English Test Band 4 (CET-4), the S-vs-M contrast remained significant for final logical score (*M*_S_ = 7.95 vs. *M*_M_ = 7.27; linear mixed model (LMM), *b* = 0.759, 95% CI [0.546, 0.972], *z* = 6.99, *P* < 0.001; Figure 3d) and recognition (*M*_S_ = 0.97 vs. *M*_M_ = 0.95; GEE binomial-logit, *b* = 0.718, OR = 2.05, 95% CI [1.45, 2.90], *P* < 0.001; Figure 3e). Both repeated-learning conditions also outperformed once-learning on final logical score (S vs. O: *b* = 2.677, *z* = 24.33, *P* < 0.001; M vs. O: *b* = 1.918, *z* = 17.82, *P* < 0.001) and recognition (S vs. O: *b* = 2.161, *P* < 0.001; M vs. O: *b* = 1.443, *P* < 0.001; Supplementary Table 4). Recognition confidence followed the same ordering: both repeated schedules exceeded once-learning (*P* < 0.001), and in the trial-level model adjusting for recognition the overall spaced-versus-massed difference was small and not significant (*b* = 0.058, 95% CI [−0.011, 0.127], *P* = 0.10), whereas among correctly recognised words spaced learning yielded higher confidence than massed learning (*b* = 0.083, 95% CI [0.024, 0.142], *P* = 0.006; Supplementary Fig. 3; Supplementary Table 4). The adjusted estimates indicate a 0.759-point advantage on the 0–10 logic scale and roughly twofold odds of a studied-item hit; descriptive paired effect sizes are reported in Supplementary Results S4. Recognition approached ceiling in the spaced condition (∼97%), with non-parametric Wilcoxon checks consistent with the parametric pattern (Supplementary Results S3; Supplementary Table 3). CET-4 moderation analyses are reported in the Supplementary Materials because they were exploratory and specific to recognition (Supplementary Table 5).

We therefore next asked whether this delayed divergence was accompanied by differences in baseline-relative retrieval-stage differentiation during the final test.

### Neural Pattern Similarity Results

#### Late NPS differentiation associated with explanatory retrieval fluency

Formal inference covered the bilateral central domain across the whole post-stimulus epoch: nine right-central and nine left-central searchlight centres at 71 window centres from 50 to 750 ms, corrected as one family per path. Figure 4c introduces the baseline-relative NPS measure: final-test neural patterns for spaced and massed items were each correlated with those for once-learning items, yielding spaced–once and massed–once NPS. Both time courses were positive at every window centre of the 50–750-ms inference epoch (Supplementary Fig. 1), indicating retained pattern correspondence with the once-learning reference. Because the same once-learning reference anchored both contrasts, the inferential question was whether spaced–once NPS differed from massed– once NPS, rather than whether either curve differed from zero.

**Figure 4.**
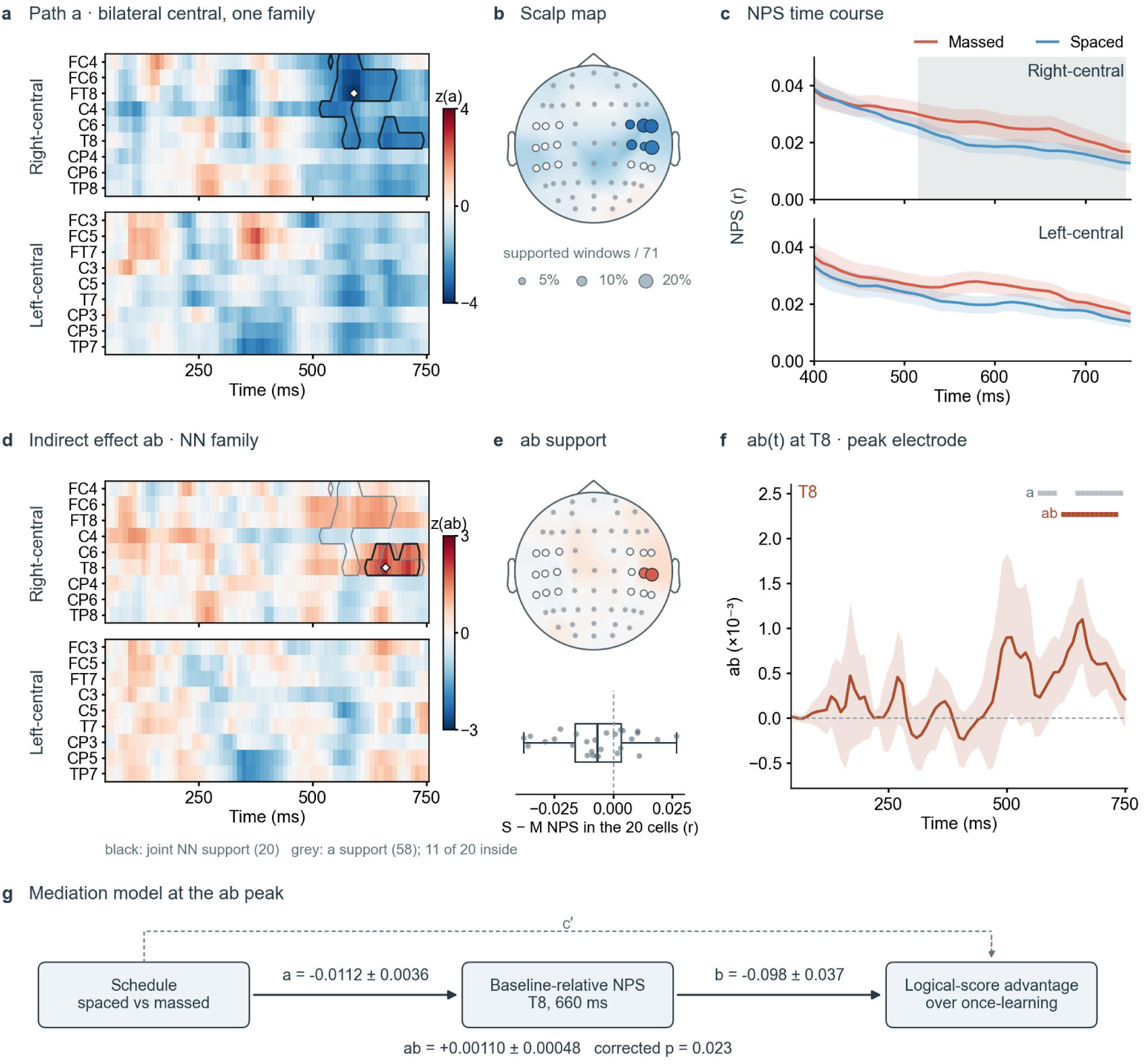
Bilateral central path-a and indirect-effect fields, their sensor support, and the mediation model at the peak cell. **a**, Signed Wald-z of path a (spaced-minus-massed difference in baseline-relative NPS) for the right-central (top) and left-central (bottom) centres at 71 window centres, 50–750 ms; blue, lower NPS under spacing (a < 0); scale ±4. Black outline, the 58 TFCE-corrected cells (one family, both hemispheres and both signs of a; Methods); white diamond, the most negative z (FT8, 590 ms). **b**, Sensor map of the a support: bubble area, supported window centres out of 71 (key: 5, 10, 20%); fill, mean signed z over those windows; open circle, tested centre without corrected support; background, all-window mean z of the 59-centre map; grey dots, other electrodes. Scalp positions, not sources. **c**, Descriptive NPS time courses: trial-mean NPS of massed (red) or spaced (blue) final-test trials to once-learning trials, averaged over each side’s nine centres; lines and bands, mean ± s.e.m. (N = 28); 400–750 ms; grey band, window centres supported in **a** (full epoch, Supplementary Fig. 1). **d**, Signed Wald-z of the indirect effect ab on the grid of **a**; red, ab > 0; scale ±3. Black outline, the 20 cells supported in the NN configuration (a < 0, b < 0); no NP (a < 0, b > 0) or left-central cell was supported. Grey outline, the a support; white diamond, the largest z(ab) (T8, 660 ms). **e**, Top, the NN support encoded as in **b**. Bottom, participant means of spaced-minus-massed NPS over the 20 NN cells (box, median and interquartile range; whiskers, range; dots, participants); descriptive. **f**, ab(t) ± model s.e. (×10⁻³) at T8 across the 71 window centres; bars, corrected support at T8 for a (grey) and NN (colour). **g**, Mediation model at the ab peak cell (T8, 660 ms): trial pairs (one target trial with one once-learning trial; 63,250 pairs from 1,842 items; 28 participants as clusters); X, learning schedule; M, the pair’s baseline-relative NPS; Y, target-minus-once-learning final-test logical score; paths a, b and c′ (dashed); estimates in Results. The same coefficients underlie **a**, **d** and **f**.

Within this domain, 58 sensor–time cells showed lower NPS for spaced than massed learning (path *a*; minimum TFCE-corrected empirical *P* = 0.022; Figure 4a; Supplementary Tables 6–7). Every corrected cell lay over the right-central sensors, and no cell survived correction in the opposite direction on either side. Corrected support opened at 520 ms over C4, extended across FC4, FC6, FT8, C6 and T8 through 600 ms, continued over FC6 and FT8 until 670–680 ms, and persisted over T8 from 650 to 740 ms, with C6 contributing again at 660–680 ms. The TFCE maximum occurred at FT8/590 ms (*a* = −0.0106, SE = 0.00274, *z* = −3.88; Figure 4a, b). The nine left-central centres were tested in the same family and yielded no corrected cell in either direction.

Within the same domain, a late right centrotemporal set of cells also showed a corrected indirect effect. Twenty cells from 620 to 730 ms survived correction for the NN configuration (*a* < 0 and *b* < 0, hence *ab* > 0; minimum TFCE-corrected empirical *P* = 0.023; Figure 4d; Supplementary Tables 6 and 8): T8 throughout 620–730 ms, with C6 contributing between 640 and 670 ms and again between 700 and 730 ms. Eleven of the twenty cells fell inside the corrected path-*a* field; the NN configuration is defined by the signs of *a* and *b* at each cell, and its family was corrected on its own, without screening by path-*a* significance. The TFCE maximum occurred at T8/660 ms (*ab* = 0.00110, SE = 0.000476, *z* = 2.31; Figure 4d, f). At T8/660 ms, a = −0.0112 (SE 0.0036) r, b = −0.098 (SE 0.037) logical-score points per unit r and c′ = +0.49 (SE 0.11), so the spaced-associated indirect effect coexisted with a direct path (partial mediation; Figure 4g). Across these cells, spaced learning was associated with lower baseline-relative NPS, and lower NPS was associated with a larger baseline-relative logical-score advantage (the target item’s final logical score minus that of its paired once-learning item). No cell survived correction for the NP configuration (*a* < 0 and *b* > 0) on either side, and no left-central cell survived correction for either configuration.

The NPS and mediation analyses linked baseline-relative retrieval-stage differentiation to the relative delayed logical-score advantage. Spacing has been proposed to allow the trace to be re-encoded and further consolidated (Benjamin & Tullis, 2010; Smith & Scarf, 2017). If so, spaced retrieval may rely more on meaning, whereas massed or once-learning retrieval may rely more on the visual form of the word. We examined the visual half of this prediction by relating final-test EEG patterns to representations from a fixed visual word-recognition model over posterior sensors (CORnet-Z; Figure 5; Methods).

**Figure 5.**
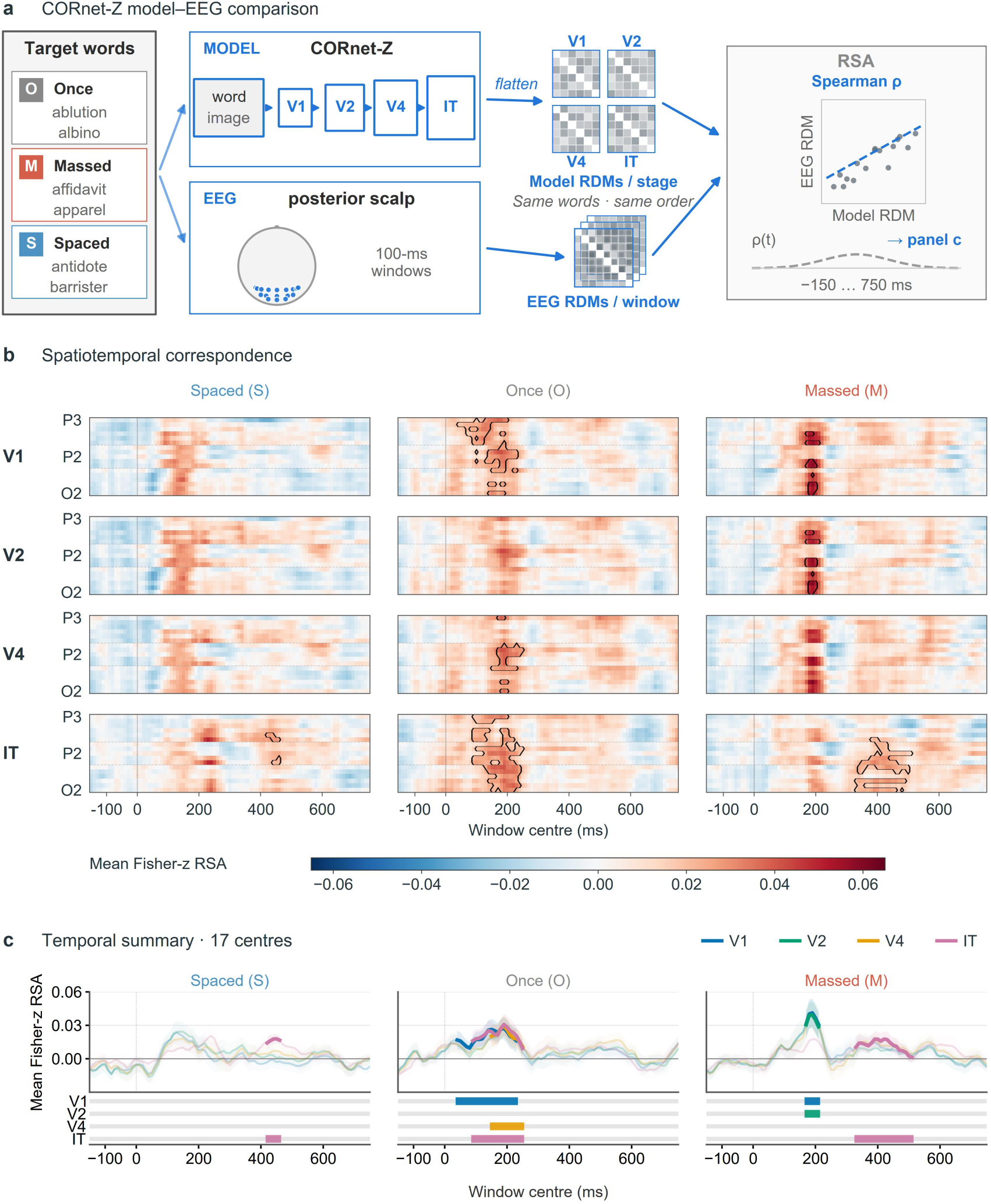
Correspondence between final-test EEG and CORnet-Z visual word representations. **a**, Analysis schematic. The 150 target words were assigned to spaced (S), once-learning (O) and massed (M) conditions (50 per condition; assignments rotated across participants). Word images yielded model representational dissimilarity matrices (RDMs) at V1, V2, V4 and IT. Representational similarity analysis (RSA) compared matching model and EEG RDM entries for the retained words within each participant and condition (Spearman correlation, Fisher-z transformed). Matrices, scatter points and the dashed curve are schematic. **b**, Participant-mean Fisher-z RSA at 17 posterior searchlight centres and 91 window centres (100-ms windows, 10-ms steps, −150 to 750 ms; N = 28). Within each stage, electrode order from top to bottom is P3, P5, P7, PO3, PO7, O1, Pz, P1, P2, POz, Oz, P4, P6, P8, PO4, PO8, O2. Black outlines mark positive RSA at family-wise corrected P < 0.05: one-sample t(27), converted to normal-equivalent Z before threshold-free cluster enhancement (TFCE; H = 2, E = 1), with 5,000 whole-participant sign flips. Correction covered each condition × stage’s 17 × 91 field separately, not the 12 families jointly. **c**, Time courses average centres within each participant, then show the participant mean ±1 s.e.m. Opaque segments and coloured bars identify windows with at least one corrected centre in b, not significance of the spatially averaged curve. Bars follow V1, V2, V4, IT from top to bottom. These exploratory tests detected IT correspondence in all three schedules; they do not establish differences between schedules.

### Model-Based Representational Similarity Results

#### Correspondence with a visual word model over posterior sensors

Final-test EEG showed corrected correspondence with CORnet-Z at one or more stages under each learning schedule (Figure 5b, c; Supplementary Table 10). Once-learning items showed support at V1 (40–230 ms), V4 (150–250 ms) and IT (90–250 ms; minimum corrected *P* = 0.006 for IT). Massed items showed early V1/V2 support (170– 210 ms) and later IT support (330–510 ms), whereas spaced items showed support only at IT (420–460 ms; minimum corrected *P* = 0.043). These intervals denote window centres with corrected support at one or more posterior centres, not support at every centre throughout the interval. IT correspondence was therefore detectable under all three schedules, while corrected early-stage correspondence was present only for once-learning and massed items. This pattern is descriptive: direct contrasts between schedules yielded no corrected cell (Supplementary Methods SM12; Supplementary Results S6).

## Discussion

Spaced learning was associated with greater late neural differentiation from the once-learning reference than massed learning, and it yielded more frequent revisions of written responses at the second initial test (T2, the test following the second learning episode, Encoding 2), despite comparable immediate logical scores under spaced and massed schedules, suggesting that the memory had changed rather than merely strengthened. The change appeared to have a direction: the mediation analysis linked lower NPS to a larger relative advantage in self-rated explanatory retrieval fluency, and in within-condition CORnet-Z tests corrected early-stage correspondence with the visual word model was detected only under massed and once-learning, whereas IT correspondence was detectable under all three schedules, although direct contrasts between schedules yielded no corrected difference. Different learners also appeared to move the same way: under spacing, revised written responses moved closer to other learners’ formulations. Together, these findings support an account in which spacing changes the organisation of meaning available for later reconstruction.

### Higher delayed explanatory retrieval after spacing despite a common mastery endpoint

To ask what remains after learners have reached a common mastery endpoint, we allowed study time to vary rather than fixing exposure duration. During Encoding 2, participants took markedly longer to reach the same criterion-verified endpoint in the spaced condition than in the massed condition (*d*_z_ = 0.96), yet the spaced condition showed no corresponding immediate logical-score advantage (*d*_z_ = 0.13). Participants in both schedules had judged the material sufficiently learned to stop studying and then met the same production criterion. Their comparable logical scores indicated similar self-rated understanding at those endpoints. The longer second-session duration was therefore the observed cost of reaching that endpoint under spacing. This pattern is consistent with desirable-difficulty accounts (Bjork & Bjork, 1992, 2020).

These similar ratings nevertheless coexisted with different rates of written-response revision. These behavioural endpoints matched immediate memory expression and self-rated coherence, but did not show that the traces supporting them were organised alike; the written records provided an observable distinction. Spaced learning produced more frequent but smaller revisions between the two initial tests, and the revised responses moved toward other learners’ formulations; massed learning produced fewer revisions, which were larger and showed no such convergence (Figure 3c, f). We read this as two modes of restructuring: after massed repetition, revision was rare but extensive and idiosyncratic; after spaced repetition, it was frequent, small, and convergent. Comparable self-rated understanding at a common mastery endpoint may therefore already have rested on differently organised memory traces (Supplementary Results S7).

Time provided the test. Both schedules entered the same retention interval from that functionally matched endpoint, so the delayed comparison asked how each organisation survived the same period of trace weakening. The traces need not have decayed at different rates: comparable weakening can disrupt a loosely integrated structure more than one whose elements are better connected (Bradshaw & Anderson, 1982; Yang et al., 2025). Two buildings can begin with comparably strong bricks and face the same weather, yet fail differently because those bricks are joined differently. The final test therefore asked which organisation better supported reconstruction after the delay.

At delayed test, both repeated schedules outperformed once-learning in self-rated explanatory retrieval fluency (Supplementary Results S4). Spacing nevertheless yielded more fluent recovery of the trained explanation than massing (LMM, *b* = 0.759, 95% CI [0.546, 0.972], *P* < 0.001). The logical score indexed participants’ perceived fluency in recovering the trained form–etymology–meaning relation during retrieval. Recognition also favoured spacing (OR = 2.05, 95% CI [1.45, 2.90], *P* < 0.001), providing convergent evidence for a broader delayed-memory advantage but not, by itself, for deeper explanatory reconstruction. Confidence followed the same pattern: among correctly recognised words, participants were less sure of massed than of spaced items (Supplementary Results S4; Supplementary Fig. 3). In retrospect, the comparable self-rated understanding that both schedules reached at the second-session endpoint reads, for massed learning, like the illusion of competence that fluent massed practice produces (Soderstrom & Bjork, 2015; Kornell & Bjork, 2008). After the delay, massed learning fell behind spacing in accuracy and in confidence alike.

### Late right-central differentiation among accessible memories under spacing

Because the neural analyses included only correctly recognised, confidence-thresholded items (Methods), the contrast was not between remembered and forgotten trials, but between organisations supporting memories that remained accessible in each schedule. Final retrieval therefore served as a common probe of organisations already established during Encoding 2 and carried through the same delay: on the present account, it did not create the schedule-linked difference but made its surviving expression observable as participants recovered the explanatory relation.

At final retrieval, spacing was associated with lower late NPS than massing, indicating greater local differentiation from the once-learning reference. Both repeated conditions retained positive correspondence with that reference at every window centre of the 50– 750-ms inference epoch (Figure 4c; Supplementary Fig. 1). The contrast thus concerns the degree of correspondence with a common reference at particular sensor–time locations. Because once-learning and the second learning episodes in both repeated conditions occurred on Day 3, all baseline-relative contrasts were observed after the same interval to the Day 5 final test.

Path-*a* identified schedule-linked baseline-relative differentiation; the separately corrected indirect effect identified where lower NPS under spacing was associated with a larger relative delayed logical-score advantage.

Xue (2022) frames retrieval as reconstruction from a progressively transformed trace; the extension here is that schedule-linked organisation matters not merely because a trace is transformed, but because its local departure from the once-learning reference can differ between schedules and only some departures remain linked to delayed explanatory retrieval.

Analysis within the bilateral central domain identified a corrected path-*a* field in which spaced learning was associated with lower baseline-relative NPS than massed learning (58 sensor–time cells; minimum TFCE-corrected empirical *P* = 0.022). The field was confined to the right-central sensors and to the second half of the epoch: it opened at 520 ms over C4, covered frontocentral and temporal sites through 600 ms, thinned to FC6, FT8, C6 and T8 by 670–680 ms, and persisted over T8 until 740 ms. The nine left-central centres entered the same correction family and showed no corrected support.

The retrieval literature offers functional and temporal benchmarks for the three overlapping portions of this field (frontocentral, frontotemporal, and T8/C6); scalp NPS indicates where and when schedule-linked differentiation is expressed, not where it is generated, and it indexes correspondence with the once-learning reference rather than item-specific reinstatement. The field opened over right frontocentral sensors (C4, FC4; 520–600 ms), a scalp distribution and onset compatible with the right central-frontal (frontocentral) old/new subcomponent that begins at 500–590 ms in word recognition (Johnson et al., 1998a) and recurs across recognition and cued recall (Friedman & Johnson, 2000). That subcomponent is usually interpreted as post-retrieval processing of recovered content; in the cited experiments it followed whichever item type required a further judgement, so it does not index retrieval success itself (Friedman & Johnson, 2000; Hayama et al., 2008). Over right frontotemporal sensors (FC6, FT8) the field was present from 550 to 680 ms. In cued recall of words, locally generated temporal activity arises first over the right hemisphere and extends furthest anteriorly there (Johnson et al., 1998b). Magnetoencephalography (MEG) source maps of delayed word recognition identify the 500–700 ms response that distinguished new from repeated words in right anteromedial temporal cortex, which the authors attribute to the semantic circuits of anterior temporal and ventral prefrontal cortex (Dhond et al., 2005), and response-locked intracranial recordings identify an anterior temporal network that reinstates conceptual content (Kragel et al., 2021). This portion of the field is compatible with the functional context of conceptual recovery described in these studies.

The latest-lasting portion lay over the T8 and C6 sensors (570–740 ms, an envelope of intermittent support), and corrected support for the indirect effect was confined to these sensors at 620–730 ms. Item-specific reinstatement in anterior temporal cortex, which represents stimuli at a conceptual level, accompanies recognition of scenes from 250 to 1150 ms (Rau et al., 2025); lateral temporal item reinstatement tracks retrieval confidence at a later, seconds-scale time point (Pacheco Estefan et al., 2019); and reinstatement of encoding patterns at 500–800 ms correlates with recognition accuracy across participants (Johnson et al., 2015). The behaviour-linked portion of the field therefore falls where the retrieval literature places the recovery of conceptual content.

The indirect effect fell within this late right centrotemporal region, concentrated over T8. Twenty cells from 620 to 730 ms survived correction for the NN configuration (minimum TFCE-corrected empirical *P* = 0.023), centred on T8 with intermittent C6 involvement. At these cells, spaced learning was associated with lower NPS, and lower NPS was associated with a larger baseline-relative logical-score advantage. Together they define a late, right-temporal locus at which greater differentiation under spacing covaried with the relative advantage in explanatory retrieval, while the direct path from schedule to that advantage remained — a partial statistical mediation. Target-selective EEG reinstatement has been reported from a similar late interval (Moccia et al., 2025).

The mediation already constrains the direction of the spaced-specific departure. Both signs are opposite to those predicted by the reinforcement account set out in the Introduction: spaced items resembled the once-learning reference less, not more, and lower rather than higher similarity accompanied better explanatory retrieval. Set beside the retrieval benchmarks above, a departure that covaries with better explanatory retrieval is compatible with reorganisation toward meaning-based structure under spacing (Boundary conditions).

### Early visual-form correspondence detected only without spacing

RSA also suggests what the mediation cannot: what the departed patterns left behind. Over posterior sensors, early-stage correspondence with the visual word model (V1, V2 or V4) survived correction only for once-learning and massed items; under spacing, no early-stage window reached correction, and correspondence with the model persisted only at IT, the stage trained to discriminate word identity. The direct contrasts for the visual word model yielded no corrected cell, so this is a dissociation in the pattern of support, not an established difference. Set beside the central-sensor result, the schedule without detectable early-stage correspondence was the one whose late right-central patterns had departed furthest from the once-learning reference and carried the behaviour-linked indirect effect. On the present account, it is as if once-learning and massed retrieval still leaned on the form of the word, whereas spaced retrieval had moved beyond it, toward its meaning. This leaves the third question: whether different learners move in the same direction. The written responses suggest that they do: under spacing, revised responses moved closer to other learners’ formulations during learning (Figure 3f), a pattern that accumulating varied contextual detail would not by itself predict.

### Reconciling encoding similarity with retrieval differentiation

On this reading, massed and spaced restudy had left differently organised traces by the end of Encoding 2 despite the same immediate mastery endpoint, and only a late subset of that difference remained linked to the delayed logical-score advantage at final retrieval. Encoding-stage studies, however, have associated spacing with greater similarity across learning encounters, not difference.

Feng et al. (2019) recorded EEG during the second presentation of novel faces repeated within a single session. Their item-specific spatiotemporal pattern similarity (STPS) across the two presentations was greater for short-spaced than massed repetitions in a late right-frontal window (606–652 ms within a 543–727-ms memory-related cluster) and partially mediated the spacing effect on subsequent recognition strength. Feng et al. interpreted this spacing-related similarity as evidence for study-phase retrieval and pattern reinstatement, rather than for greater encoding variability.

Zou et al. (2025) used 7 T fMRI with continuous recognition of natural scenes repeated at lags from seconds to months. Among encounters correctly recognised at E2, stimulus-specific E1–E2 similarity increased with lag in ventromedial prefrontal cortex (vmPFC) and was positively related to recognition at the third presentation (E3) only at longer lags, whereas similarity in early visual cortex decreased with lag. The opposing regional directions in Zou et al. also caution against treating E1–E2 similarity at Encoding 2 as a unitary index of reinstatement.

Both studies therefore estimated item-specific similarity across two learning encounters. In contrast, our NPS reads each repeated-learning condition at final retrieval against a once-learning reference whose sole study episode was aligned with the second learning episodes on Day 3. On the surface, these results point in opposite directions: prior work associated spacing with greater item-specific similarity across learning encounters, whereas our late retrieval analysis identified a larger local departure from the once-learning reference under spacing than under massing. That distinction alone does not resolve the tension. How can spacing be associated with greater continuity across learning encounters yet with greater differentiation from the once-learning reference at final retrieval?

Separating the observation window from the similarity reference clarifies what E1–E2 similarity can and cannot tell us. Greater E1–E2 similarity describes continuity across the two learning encounters; it does not by itself reveal how much of the first learning episode (Encoding 1) remained uninternalised and directly available to shape reconstruction during Encoding 2, versus how much had already been assimilated into the structure guiding that reconstruction.

We propose that Encoding 1 can shape Encoding 2 through two coexisting forms of influence: an uninternalised, episode-bound product that remains directly available, and an influence already assimilated into the knowledge structure that organises reprocessing. Hanfstingl et al. (2021) characterise assimilation as the more economical, knowledge-driven route when incoming information can be fitted to existing structure. Building on this distinction, we propose that spacing changes the relative weight of these forms rather than whether Encoding 1 influences Encoding 2.

After a short massed interval, a larger episode-bound remainder may remain available during Encoding 2, allowing familiarity and direct comparison to contribute while the prior representation remains relatively accessible. Consistent with the limited accessibility component of this proposal, McKinley et al. (2019) found that recently encountered related information reduced subsequent self-paced study time at short lags, with the relatedness effect observed under massed conditions. With spacing, a greater share of the first encounter may have been assimilated into the learner’s knowledge structure, shifting the balance toward the assimilated constraint while leaving the uninternalised remainder available but relatively less influential during Encoding 2 (Figure 6). Assimilation need not substantially alter an established schema, especially for material that does not challenge prior knowledge, such as the faces used by Feng et al. (2019). The same input may then be processed through a largely stable system, producing greater E1–E2 similarity even while Encoding 2 reorganises the trace more substantially.

**Figure 6.**
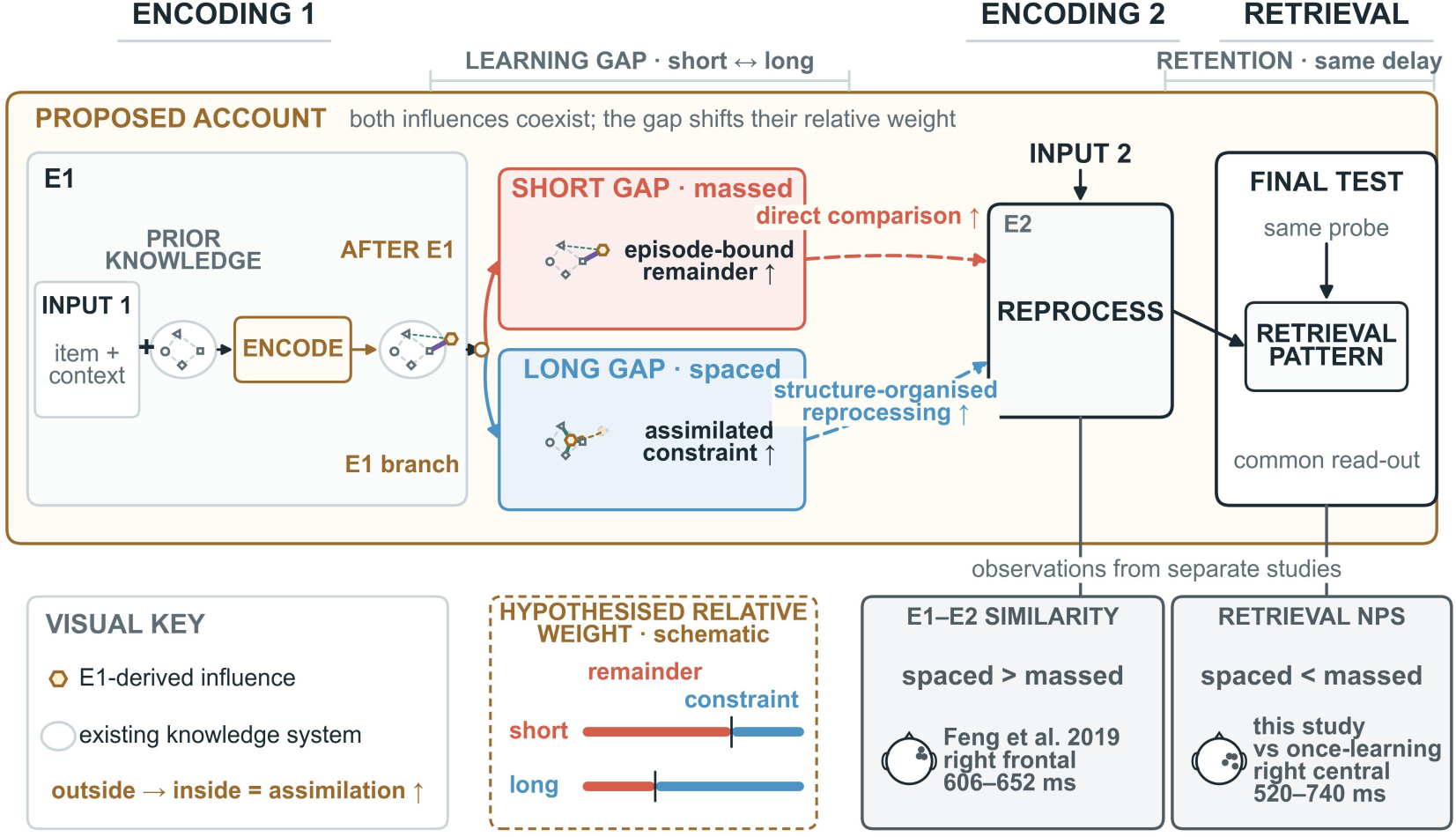
A proposed account reconciling encoding-stage similarity with retrieval-stage differentiation. In the proposed account (boxed region), Encoding 1 leaves an uninternalised, episode-bound remainder and an influence already assimilated into the knowledge structure; both remain available during Encoding 2. A short (massed) gap is proposed to favour the remainder and direct comparison, whereas a long (spaced) gap favours the assimilated constraint, which organises reprocessing; in the present design the final test applies the same probe to both schedules after a common delay. Lower cards (observations from two separate studies, placed at their measurement stages, not a demonstrated transition): left, greater item-specific E1–E2 pattern similarity for short-spaced than massed repetitions over right-frontal sensors at 606–652 ms during the second encoding (Feng et al., 2019); right, lower retrieval-stage NPS to the once-learning reference for spaced than massed items over right-central sensors at 520–740 ms in the present study (Figure 4a). The relative weights, the assimilation transition and the two proposed contributions to reprocessing are hypothesised and were not measured. Bar lengths in the key illustrate the proposed shift; they do not specify an exhaustive partition or estimated weights. Scalp positions summarise sensor-space support, not sources.

These are hypotheses about how the schedule-dependent organisations were established during Encoding 2. The present final-retrieval EEG does not observe those learning-stage operations; it reads out the resulting organisation at final retrieval. Zou et al.’s phrase, “re-encoding of past experience,” captures the logic of this proposal: the influence of Encoding 1 is neither absent from Encoding 2 nor reducible to replay.

The first encounter becomes less an *endpoint* to be recovered than an *entry condition* for reconstruction. The encoding-stage and retrieval-stage findings are best read as successive observation windows on repeated learning: earlier studies sampled the process-mixed re-encounter itself, whereas the present final test sampled the schedule-linked organisation that survived the common delay. Reconstructing the explanatory relation at final test was the common probe of that surviving organisation, not the operation that created it.

### Boundary conditions and future tests

The first boundary concerns the type of learning. The etymological relations used here likely place the task closer to the assimilative than the accommodative end of meaningful learning: they can be incorporated into an existing lexical–semantic system without requiring fundamental revision of that system. These data therefore do not establish whether the proposed weighting account would hold when learning requires accommodation, as in misconception correction or conceptual change. In such cases, Encoding 1 may alter the organising schema enough that spacing need not increase E1– E2 similarity, even if delayed reconstruction still improves.

The delayed behavioural endpoint sets another boundary. The final logical score was a task-trained, participant-rated index of perceived explanatory retrieval fluency. Participants rated how completely and fluently they had reconstructed the etymological chain, but no written or spoken explanation was externally scored. The score therefore supports a claim about perceived explanatory retrieval fluency, not objective explanatory accuracy. It cannot by itself distinguish whether spacing improved the recovered relation, made that relation easier to access once the word itself was recognised, or changed the metacognitive appraisal of that access. Future work should collect written or spoken reconstructions, score individual relational links blind to condition, and test whether late retrieval-stage NPS differentiation predicts objective explanatory completeness beyond recognition, confidence, and self-rated fluency.

A separate boundary concerns the neural interpretation. The mediation effects link this retrieval-stage read-out to the logical-score advantage. Condition specificity, temporal organisation, and behavioural linkage support describing the lower-NPS field as structured differentiation. It does not exclude the possibility that repeated study merely made each word’s own pattern more distinct without changing its organisation, which could also lower similarity across different words, and indirect effects in which spacing raised similarity were not tested. The causal direction of this association remains to be tested experimentally. A stronger test would alter the relative accessibility of the uninternalised, episode-bound Encoding 1 product at Encoding 2—for example, through retrieval support or intervening interference—while holding the immediate mastery endpoint and final retention interval constant. If the weighting account is correct, changing that accessibility should shift the balance between episode-bound and assimilated influences and thereby alter both late retrieval-stage NPS differentiation and objectively scored explanatory reconstruction. A further caveat concerns trial selection. Because repeated learning usually left more remembered items than once-learning, massed and spaced trials were ranked by final logical score and the highest-scoring were retained up to the once-learning count (Methods), so that the schedules were compared on equal numbers of well-reconstructed items rather than on sets differing in size and quality; the effect of this selection on the indirect effect was not assessed.

The representational analyses draw a boundary of their own. They say which fixed model geometry the final-test patterns resembled, condition by condition; their domains and layers were fixed only after earlier scans had been inspected, so those results are exploratory; and they do not test whether the mediating pattern is itself semantic, nor compare hemispheres. The right-central support and the null left-central result sit in one correction family, so lateralisation would need a direct hemisphere × schedule test. That test, and a pre-specified conjunction of the NPS and RSA supports, are the natural next steps for lateralisation and overlap; identifying semantic content in the mediating pattern would require a model-based test of that pattern itself.

A final design boundary concerns what this comparison estimates. Criterion-verified mastery, exposure time, and temporal order could not be fixed independently. We therefore prioritised a common mastery endpoint and aligned each condition’s final or sole learning episode on Day 3, so all conditions entered the Day 5 test after the same retention interval; first-session study duration and first-session logical score were covariate-adjusted. The resulting contrast is a condition-level comparison of complete, mastery-gated schedules, not a pure interval effect with all schedule-contingent practice and strategy development held constant.

## Conclusion

Spaced and massed learning reached the same criterion-verified mastery endpoint with comparable immediate ratings of understanding, yet spacing yielded higher self-rated explanatory retrieval fluency after the common Day 3–Day 5 delay. This advantage was accompanied by late right-central differentiation from the once-learning reference, with a partly overlapping indirect-effect pattern over T8 and C6 linking lower NPS to the relative logical-score advantage. Together with written revisions that drew closer to other learners’ formulations, this pattern suggests that spacing changes the memory trace rather than merely strengthening it, in a direction linked to self-rated understanding. These findings are consistent with the view that learners can each regard an explanation as coherent and understood, yet differ in the degree to which they have reconstructed it. Such felt completeness marks a provisional endpoint within the learner’s current knowledge, one that spacing appears to move, not a fixed ceiling on understanding.

## Methods

### Participants

A sensitivity calculation indicated that 24 participants would give 80% power for a medium within-participant effect (Supplementary Results S5). Participants were a nonprobability (convenience) sample of undergraduate students from a university in China. After exclusions made before analysis on EEG quality or on laboratory records of the sessions (Supplementary Methods SM1), the final sample comprised 28 participants (21 female, 7 male; mean age 19.7 ± 0.9 years, range 19–21, estimated from year of enrolment). For the written-response revision measure, two further participants were set aside because fewer than 20 of their words met the both-responses-observed criterion in at least one condition, leaving 26; this coverage rule was applied to the written records alone, without reference to any outcome. The sample was predominantly female, and future mixed-gender studies are recommended to assess whether the observed behavioural and neural effects generalise across sexes. All participants had normal or corrected-to-normal vision and were right-handed. Reported College English Test Band 4 (CET-4) scores averaged 462.8 ± 32.3 out of 710 (range 400–542). The study was approved by the Institutional Review Board of Chongqing University of Arts and Sciences, where the data were collected, and was conducted in accordance with the Declaration of Helsinki. Participants provided written informed consent following explanation of EEG procedures and voluntary participation principles. They received 150 yuan (∼20 USD) compensation upon completion.

### Stimuli

Learning items were English–Chinese word pairs chosen to be unknown to our participants yet decomposable into words they already knew. Candidate words were drawn from GRE vocabulary lists with CET-4 words removed, so that they lay above the participants’ level (CET-4 scores 400–542), and were pre-rated for familiarity by 27 undergraduates from the same population who did not take part in the experiment; the 150 least familiar words were retained.

Each item was presented with its Chinese translation, familiar component words belonging to CET-4 vocabulary, and an etymological explanation linking those components to the target meaning, so that the meaning could in principle be derived from known parts (examples are provided in Supplementary Table 9). Word-set construction, counterbalancing of sets across conditions and the 100 novel foils used in the recognition test are described in Supplementary Methods SM2.

### Experimental Design and Procedure

The experiment was programmed in MATLAB (R2019a) using Psychtoolbox (Brainard, 1997; Pelli, 1997) and conducted in a quiet, soundproof laboratory with controlled lighting. Participants sat 60 cm from a 17-inch LCD monitor (1400×900 resolution, 60 Hz refresh rate).

#### Practice Phase (Day 0)

Participants practised with 20 words to become familiar with Grimm’s laws, the structure of the learning materials, and the experimental procedure. They continued until they achieved 100% accuracy on a vocabulary sheet, using a new practice set when necessary. This practice also introduced the difficulty of the subsequent learning task.

#### Learning Phase (Days 1 and 3)

As illustrated in Figure 1b, each 25-word set was presented sequentially. Each trial began with a 2-s cue instructing careful reading, after which the English word appeared alone until participants pressed a key, during which they attempted to recall its meaning (self-paced). The word’s Chinese definition and etymological explanation were then presented for self-paced study. Participants used the etymological relation to memorise the word and rated it as “confident”, “unsure” or “again“; only words rated “confident” were removed from the current study cycle. Once all 25 words had been rated “confident”, participants completed an initial test in which, for each English target, they wrote the intermediary words and Chinese definition and rated the logic of their reconstruction (the first- and second-session logical scores). A score of 100% was required; otherwise, the set entered another study–test cycle.

The interval between the two learning episodes was approximately 48 h in the spaced condition and approximately 2 h in the massed condition (Figure 1a). On Day 1, participants completed the first learning episode for the two spaced sets. On Day 3, they completed the first learning episode for the massed and once-learning sets, followed by the second learning episodes for the spaced and massed sets. Consequently, the second learning episodes for both repeated conditions and the sole scheduled learning episode for the once-learning condition occurred on Day 3, providing the same planned interval to the Day 5 final test. Two participants deviated from this calendar (one completed the sessions on Days 1, 2 and 4, the other on Days 1, 5 and 9). Because every participant completed the once-learning episode and the second learning episodes of both repeated conditions in a single session, the interval to the final test was nevertheless identical across conditions within each participant (36.5–52.1 h for 27 participants and 85.9 h for one). The initial test after each set required the same 100% production criterion across conditions.

#### Final Test (Day 5)

As shown in Figure 1c, each trial began with a 1-s fixation cross, followed by a 2-s presentation of an English word (height: 0.7 cm, 0.67° visual angle; width: 2–3 cm, 1.9°–2.9°). During the word presentation, participants attempted to retrieve its learned meaning and reconstruct the etymological relation linking the English form to the Chinese meaning—the same operation practised during the initial tests. After the word disappeared, participants made an old–new recognition judgement (4-s response limit), with 100 novel foils intermixed, rated their recognition confidence from 0 (guess) to 5 (very confident), and rated their explanatory retrieval fluency on a 0–10 scale (final logical score, below). The Day 0 words were used for practice, and the instructions were restated before testing. A self-paced break was offered after every 50 trials; the final test lasted approximately 35–40 min.

#### Temporal structure and matched endpoints

A cross-day comparison of spaced, massed, and once-learning conditions at a common final test necessarily imposes temporal ordering: when the interval from the final or sole scheduled learning episode to test is held constant, the first spaced episode must precede the Day-3 massed and once-learning blocks. This ordering follows jointly from the spacing interval and the common-retention-interval control, rather than from arbitrary block placement. We therefore aligned the final learning exposure for both repeated conditions (Encoding 2) with the sole scheduled learning episode on Day 3. By design, this held the planned interval from the final or sole scheduled learning episode to the Day 5 test at approximately 48 h in every condition (observed intervals are given above), separating differences in learning history from unequal time-to-test.

#### Condition-level comparison and alternative designs

Because all three conditions were tested within the same participant and final-test EEG recording, the condition contrasts held participant, electrode montage, and retrieval context constant (Supplementary Methods SM5). A two-group between-participant comparison would remove direct carryover between spaced and massed blocks, but it would not separate the spacing interval from schedule-contingent practice and strategy development. Adding a third once-learning group would provide the one-scheduled-episode reference required for baseline-relative NPS, but it would not solve that decomposition, because once, massed, and spaced learning necessarily entail different encounter histories. Only extensive pretraining to a common strategy asymptote could further reduce this source of variation, and that would change the question to spacing after task-specific training. Our design instead maximally matched the observable learning and test states: every condition reached the same criterion-verified endpoint at its final or sole Day-3 learning episode and was probed in the same Day-5 test. The estimand is therefore the condition-level effect of the complete mastery-gated schedule, not an item-level effect or a pure temporal-lag effect with all schedule-contingent adaptation removed (Supplementary Methods SM3).

#### Measured sequence differences

First-session study duration and first-session logical score were included as covariates in the trial-level models to adjust measured differences potentially related to task familiarisation and strategy development. The systematic study-duration pattern across conditions (S > M > O) and the first-session logical-score comparisons are reported in Supplementary Results S2; the spacing effects remained robust after covariate adjustment (see Results and Supplementary Results S4). These models improve estimation and test the robustness of the condition-level effect; they do not eliminate all unmeasured consequences of fixed order or shift the theoretical inference to individual items.

##### Logic Fluency Rating

The final-test logical score was a participant-rated measure of explanatory retrieval fluency. During learning and immediate tests, participants repeatedly practised the same operation—decomposing the English word, reconstructing the etymological link, and using that link to infer the Chinese meaning—so the final-test rating targeted the fluency of a practised operation rather than a novel introspective judgement.

At final test, after the old–new judgement and confidence rating, participants rated how fluently they had reconstructed this etymological logic during the word presentation on a 0–10 scale (0 = no relevant logic recalled; 10 = complete and fluent reconstruction of the logic chain). The measure targeted a different judgement from confidence: confidence assessed certainty that an item was old, whereas logical score assessed the perceived completeness of the reasoning chain. Because it was self-rated rather than externally scored, we treat it as a task-trained subjective index of explanatory retrieval fluency and as a proxy for constructive understanding, not as an externally scored objective measure.

### Behavioural Data Analysis

#### Behavioural outcomes

Three outcome measures were assessed at final test: studied-item recognition accuracy (hit rate; the proportion of learned words correctly judged “old”), final logical score rating (a task-trained self-rated index of explanatory retrieval fluency based on the perceived completeness of etymological reasoning), and confidence ratings. Because the condition-wise behavioural dataset contained only studied items, this recognition measure indexes hit rate rather than old–new discrimination; responses to novel foils were not included in these models. In addition, a learning-phase process measure, the written-response revision rate between the first and second initial tests (T1 and T2; T2 is the Encoding 2 test), was defined for the two repeated conditions as the proportion of a participant’s words for which the normalised written response (intermediary words plus Chinese definition) at the second initial test differed from that at the first; a word entered the denominator only if both responses were observed. For words whose response changed, two further quantities were derived from text embeddings (BGE-M3; Chen et al., 2024): each response was represented as the joint string of intermediary words and Chinese definition (Unicode-normalised) and encoded as the model’s L2-normalised [CLS]-token embedding. Revision size was the cosine distance between a word’s T1 and T2 embeddings. To index how close a response was to other learners’ formulations, the group centre for a word at a given test was the normalised mean embedding of all other participants’ responses to that word at the same test (leave-one-out, computed only when at least five other responses were available). Distance to the group centre was the cosine distance between the participant’s response and that centre, taken at T1 against the T1 centre and at T2 against the T2 centre. Its T1-to-T2 change (Δ) served as the convergence outcome (Figure 3f; both pipelines are illustrated in Supplementary Fig. 2).

#### Omnibus tests

Because several measures deviated from normality—right-skewed study durations and a ceiling-related deviation in studied-item hit rate (≈97% in the spaced condition; Supplementary Results S1; Supplementary Table 2)—we used Friedman tests, a non-parametric omnibus suitable for within-participant designs, for all comparisons across the three conditions (spaced, massed, once-learning). The same test was applied uniformly, including to approximately normal measures such as first-session logical score, for consistency.

#### Pairwise comparisons

Post-hoc paired comparisons used paired *t*-tests with Holm correction, with effect sizes reported as Cohen’s *d*_z_ (Lakens, 2013; Supplementary Methods SM4). For the studied-item hit rate, whose deviation from normality was ceiling-related, Wilcoxon signed-rank tests were additionally conducted and yielded the same conclusions (Supplementary Results S3, Supplementary Table 3).

#### Trial-level regression models

To control for baseline differences in first-session logical score and study duration across conditions, trial-level outcomes were analysed while accounting for repeated observations within participants. Linear mixed-effects models (LMMs) with participant-specific random intercepts were used for final logical score and confidence; the confidence models additionally included the studied-item hit outcome and its interaction with condition; the confidence contrast is reported with the hit outcome grand-mean centred and, in a model without the hit term, for correctly recognised words only (Supplementary Results S4). The binary studied-item hit outcome was analysed using generalised estimating equations (GEE; Liang & Zeger, 1986) with a binomial distribution and logit link, participant as the clustering variable, and an exchangeable working correlation structure. Reported coefficients come from the full model, which included condition, the first-session covariates, and CET-4 and its interaction with condition; the stepwise build-up is given in Supplementary Results S4, with full coefficients in Supplementary Table 4.

#### Word-level mixed models for written-response revision

Revision outcomes were analysed at the word level with crossed random intercepts for participants and words (lme4 1.1-35.5, Bates et al., 2015; lmerTest 3.1-3, Kuznetsova et al., 2017; R 4.4.2). Whether a word was revised (0/1; 2,550 observable words from 26 participants) was modelled with a logistic generalised linear mixed model; revision size and Δ distance to the group centre (760 revised words) with linear mixed models (restricted maximum likelihood, REML; Satterthwaite degrees of freedom). Fixed effects were condition (spaced vs massed), CET-4 score and its interaction with condition, first-session study duration and first-session logical score, each grand-mean centred over the 2,550 observable words (the convergence model re-centred over its 760 revised words). The reported condition effect is the spaced-minus-massed difference at that grand mean of CET-4 (an odds ratio for revision, otherwise on the cosine-distance scale), with Wald 95% confidence intervals. Participant-only random-intercept models, an alternative covariate convention, and a fixed-centre variant of the convergence measure are reported in Supplementary Results S7.

#### Exploratory moderation analysis

English proficiency (CET-4 score) was examined as an exploratory moderator of the spacing effect through its interaction with condition; because the moderation was specific to recognition, it is reported in the Supplementary Materials (Supplementary Table 5).

### EEG Recording

EEG signals were recorded using a 64-channel electrode cap (Brain Products actiCHamp system, 2015) based on the extended international 10-20 system. Electrode impedances checked before recording had a median of 3 kΩ, with 88% at or below 5 kΩ and all but one at or below 10 kΩ (logged for 27 of the 28 participants). The online reference was FCz. The ground electrode was placed at AFz. Vertical electrooculography (VEOG) was recorded from an electrode 10 mm below the right eye; horizontal EOG was not recorded. Signals were recorded at 500 Hz with an online high-pass time constant of 10 s (approximately 0.016 Hz) and no online notch filter.

### EEG Preprocessing and Trial Selection

Bad channels were identified by visual inspection of signal quality and interpolated using spherical splines (Perrin et al., 1989). The continuous data were then band-pass filtered at 0.1–30 Hz, notch-filtered at 50, 100, 150 and 200 Hz, and re-referenced offline to the average of all EEG electrodes. Independent component analysis (ICA) implemented in MNE-Python (Gramfort et al., 2013) was fitted to a copy of these data additionally high-pass filtered at 1 Hz. MNE’s find_bads_eog function (threshold = 3) provided candidate EOG components as an aid; the ICA components were then inspected manually, and components judged to reflect ocular activity were removed from the 1-Hz high-passed copy, which formed the analysed signal (1–30 Hz). The cleaned continuous data were segmented into epochs time-locked to stimulus onset (−200 to 800 ms), with the pre-stimulus interval (−200 to 0 ms) serving as the baseline (mean subtraction). Epochs with remaining artefacts were automatically repaired or rejected using the AutoReject algorithm (Jas et al., 2017; Supplementary Methods SM6).

#### Trial selection

The EEG epoch was time-locked to the 2-s word presentation, when participants attempted to retrieve the learned meaning and reconstruct the etymological logic. For the main neural analyses, trials were then filtered using the subsequent behavioural judgements: included trials required correct “old” judgements for studied items and confidence ratings ≥3 (0–5 scale). This selection ensured that the analysed retrieval attempts came from trials that participants later recognised with at least moderate confidence. Because once-learning usually yielded the fewest high-confidence correct trials, its complete surviving set defined the participant-specific target count.

Massed and spaced trials meeting the same recognition and confidence criteria were then ranked by final logical score, and the highest-scoring trials were retained up to that count; a condition with fewer eligible trials kept all of them. The comparison therefore used equal numbers of successfully retrieved trials in 24 of the 28 participants (the remaining four differed by 1–10 trials) and prioritised the strongest explanatory reconstructions available after repeated learning. After preprocessing and trial selection, each participant contributed an average of 33.0 trials per condition (range: 22–46 trials; Supplementary Methods SM7).

### Neural Pattern Similarity and Mediation Analysis

Figure 2 summarises the neural pattern similarity (NPS) analytic pipeline, progressing from searchlight-based similarity estimation (a) to two-level mediation modelling (b) and TFCE permutation inference over the bilateral central domain (c).

#### Spatiotemporal Searchlight NPS Computation

NPS was computed using a spatiotemporal searchlight approach (Kriegeskorte et al., 2006; Figure 2a). In the spatial domain, we defined 59 searchlight patches, each centred on one EEG electrode and encompassing all neighbouring electrodes within a 5-cm radius. Temporal pole electrodes (FT9, FT10, TP9, TP10) were excluded due to artefact susceptibility. In the temporal domain, we applied a sliding window of 100 ms width with a step size of 10 ms, yielding 91 overlapping time windows spanning the full epoch (parameter justification in Supplementary Methods SM8). The complete 59 × 91 maps were retained to describe the full-epoch spatiotemporal pattern; formal mediation inference was restricted to the bilateral central domain described below (Supplementary Methods SM9).

For each spatiotemporal unit (patch × time window), we extracted the multivariate EEG pattern (channels × time points) for each trial during the final test phase. Trials were grouped by their original learning condition: once-learning (O), massed (M), and spaced (S). NPS was computed as the trial-by-trial Pearson correlation between EEG pattern vectors of items from different learning conditions (Haxby et al., 2001; Kriegeskorte et al., 2008). Specifically, we computed pairwise correlations between all once-learning trials and all massed trials (O–M trial pairs), and between all once-learning trials and all spaced trials (O–S trial pairs). NPS was defined as *r* (Pearson correlation coefficient), where higher values indicate greater pattern similarity to the once-learning baseline. This procedure yielded a 59 (spatial patches) × 91 (time windows) NPS value for each trial pair, which was then entered into the multilevel mediation model.

Because NPS was computed between once-learning trials and repeated-learning trials, lower O–S than O–M similarity indicates that spaced learning produced retrieval-stage patterns that were more differentiated from the once-learning baseline. The once-learning condition therefore served as a one-scheduled-episode baseline against which the two repeated-learning schedules were compared. This baseline is itself a learned trace observed at retrieval, so departures from it index reorganisation relative to one learning episode rather than distance from a neutral zero. We interpret lower baseline-relative NPS as greater retrieval-stage differentiation from this baseline, not as direct within-condition trial-to-trial variability. Importantly, lower NPS is not inherently beneficial: its interpretation as structured differentiation depends on its condition specificity, organised spatiotemporal profile, and statistical association with logical gains.

#### Multilevel Mediation Model

To test whether condition-related differences in NPS during the final test were statistically associated with the relative delayed advantage in self-rated explanatory retrieval fluency, we fitted a two-level mediation model (Preacher et al., 2010) at each spatiotemporal unit. Models were fitted in Mplus (version 8.3; Muthén & Muthén, 1998–2017; Figure 2b) with robust maximum likelihood (MLR) on NPS expressed on its original, unrescaled Pearson-*r* scale. The input comprised 63,294 trial-pair rows from 28 participants; the 44 rows of one massed target lacking a first-session logical score were excluded listwise, so each model was fitted to 63,250 rows. Trial-pair rows formed the within-participant observations and participants formed the between level; the 1,843 participant-by-target blocks were used only to preserve intact dependency units during permutation and did not constitute a third model level. Both the mediator and the outcome were defined relative to the once-learning baseline: for each spaced or massed target item, NPS was its similarity to the paired baseline items, and the outcome was its final-test logical advantage over those baseline items. At the within level, path *a* linked learning condition (coded 1 = massed, 2 = spaced) to NPS, path *b* linked NPS to this baseline-relative logical advantage, and path *c′* represented the direct effect of condition on that advantage. NPS, first-session logical score, and first-session study duration were grand-mean centred; the two behavioural covariates were included in both the mediator and outcome equations. At the between level, random intercepts for NPS and the outcome were clustered by participant. The indirect effect was computed as *ab* = *a* × *b* and was interpreted as a statistical association under the fitted model, not as evidence of causal neural mediation (Supplementary Methods SM9).

The model was estimated across all 5,369 spatiotemporal units (59 patches × 91 time windows) to retain descriptive full-epoch maps of the *a*, *b*, *c′*, and *ab* paths; 1,218 of these units carry a fixed numerical mask (1,150 of them before 50 ms) and yield no usable estimate. Formal inference covered a bilateral central domain of 18 searchlight centres — FC3, FC5, FT7, C3, C5, T7, CP3, CP5 and TP7 over the left hemisphere and FC4, FC6, FT8, C4, C6, T8, CP4, CP6 and TP8 over the right — at all 71 post-stimulus window centres from 50 to 750 ms (18 × 71 = 1,278 units). The domain operationalised the spatial hypothesis stated in the Introduction and was fixed before any inferential map was inspected (Kriegeskorte et al., 2009): its centres span the fronto-central, temporal and centro-parietal rows of each hemisphere, and their 5-cm searchlight neighbourhoods extend from the lateral frontal to the parietal row (40 of the 59 electrodes in total, with no midline electrode). Each estimate summarised a 100-ms window indexed by its centre. The complete observed map (permutation 0) and each of the 1,991 null maps contained all 1,278 units, and TFCE was computed across this complete field with no numerically masked units.

#### TFCE Permutation Testing over the Bilateral Central Domain

To control family-wise error across the bilateral central sensor–time field, we applied threshold-free cluster enhancement (TFCE; Smith & Nichols, 2009) to the directional *z*-statistic maps using *H* = 2 and *E* = 1 (Figure 2c). Spatial adjacency was defined from electrode positions by Delaunay triangulation and restricted to the 18 selected centres, which yielded no adjacency between hemispheres; temporal adjacency was defined by successive 10-ms windows. For path *a*, the tested directions were *a* < 0 and *a* > 0. For the indirect effect, the tested directions were NN (*a* < 0, *b* < 0, *ab* > 0) and NP (*a* < 0, *b* > 0, *ab* < 0). We generated 1,991 null permutations by shuffling spaced/massed labels within each participant at the intact target-block level, preserving each participant’s condition counts and keeping all once-learning pair rows for a target together. The full MLR model was re-fitted at every eligible unit on every permutation. For each permutation, the larger raw maximum TFCE value across the two path-*a* directions formed the shared path-*a* null, and the larger raw maximum across the NN and NP directions formed the separate shared path-*ab* null. Path-*ab* inference covered the full tested domain without screening by path-*a* significance; NN and NP shared one correction family. Corrected pointwise *P* values were calculated as [1 + number of shared-family null maxima at least as large as the observed point TFCE]/1,992, with *P* ≤ 0.05 defining significance. Thus, significance was calibrated by spaced/massed condition-label permutation and shared raw-max TFCE correction; path *a* and path *ab* were treated as distinct correction families (Maris & Oostenveld, 2007; Supplementary Methods SM10).

### Model-Based Representational Similarity Analysis

Representational similarity analysis (RSA; Kriegeskorte et al., 2008) compared the geometry of the final-test EEG patterns with that of a fixed visual word-recognition model using representational dissimilarity matrices (RDMs). The model was the literate nonbiased-words CORnet-Z checkpoint (Agrawal & Dehaene, 2024). It received each of the 150 target words as an image—black 32-px Arial on a 224 × 224 canvas, with the background darkened to a grey-screen proxy and ImageNet normalisation—and returned one activation vector per word at each of its four stages (V1, V2, V4 and IT). We used IT as a high-level visual–orthographic representation supporting written-word identification (Hannagan et al., 2021; Agrawal & Dehaene, 2024); stage activations, rather than classifier outputs, were used to construct the model RDMs.

Model RDMs contained the correlation distance between the CORnet-Z activation vectors for every pair of words, with one RDM per stage. EEG RDMs used 5-cm searchlight neighbourhoods and 91 half-open 100-ms windows (10-ms step; window centres from −150 to 750 ms). The CORnet-Z analysis used the archived NPS-selected trials. For each participant, condition, searchlight and window, EEG dissimilarity was the correlation distance between the channels × samples patterns of each pair of retained trials. The upper triangle of each EEG RDM, excluding the diagonal, was compared by Spearman correlation with the model RDM restricted to the same words in the same order. Correlations were Fisher-z transformed.

Inference asked, separately for each learning condition and CORnet-Z stage, whether participant-mean Fisher-z RSA exceeded zero within the posterior domain. At each centre–window cell, the one-sample t statistic (df = 27; N = 28) was converted to a signed normal-equivalent Z before positive-tail threshold-free cluster enhancement (TFCE; H = 2, E = 1). Spatial adjacency was inherited from the full-scalp Delaunay graph and restricted to the 17 posterior centres; temporal adjacency linked successive 10-ms window centres. Each of the 12 condition × stage families comprised the complete 17 × 91 field from −150 to 750 ms. Family-wise error was controlled separately within each family using 5,000 sign flips, with the same sign applied to every cell of a participant’s map. Corrected P values were (1 + k)/5,001, where k counts null maximum TFCE values at least as large as the observed cell’s TFCE value (*P* < 0.05). No correction was applied across the 12 families, and these single-condition tests did not themselves compare learning schedules.

Comparisons between schedules in the main text are descriptive. Direct contrasts between schedules were run for CORnet-Z and are reported in the Supplement (Supplementary Methods SM12; Supplementary Results S6). The CORnet-Z RSA analysis was exploratory: the posterior sensor domain was fixed after inspection of earlier whole-scalp results. The directional prediction that spaced retrieval would lean less on the visual form of the word provides the theoretical motivation, but this data-informed domain selection means that the analysis is not reported as a pre-specified confirmatory test (Kriegeskorte et al., 2009).

### General Statistical Conventions

#### Visualisation

Complete 59 × 91 NPS maps and grand-average spaced–once and massed–once time courses were retained for descriptive full-epoch orientation. Formal inferential displays covered the bilateral central domain across 50–750 ms: complete signed Wald-*z* fields for path *a* and the indirect effect were shown for all 1,278 cells, with outlines around cells surviving family-specific shared raw-maximum TFCE correction and diamonds at the raw-TFCE peaks. Sensor maps summarise, per centre, the number of supported window centres and the mean signed *z* over those windows; they are sensor-space summaries and do not imply source localisation.

### Software and Implementation

EEG processing used MNE-Python (custom searchlight functions); trial-level LMM and GEE models used Python (scipy, statsmodels); word-level mixed models for written-response revision used lme4 and lmerTest in R 4.4.2; multilevel mediation used Mplus (Muthén & Muthén, 1998–2017), called from R via MplusAutomation with parallelisation; and visualisations used matplotlib. Software versions are listed in Supplementary Methods SM11.

## Use of AI Tools

Large language model assistants (Claude, Anthropic; ChatGPT and Codex, OpenAI) were used during data analysis and manuscript preparation, after data collection was complete, to help write and check analysis and figure code, to suggest and cross-check statistical analyses, to search and screen the literature, and to draft and edit text. The authors made all analytical and interpretive decisions, verified all outputs against the data and the cited sources, and take full responsibility for the content.

## Data availability

The de-identified behavioural data and the processed EEG data (NPS-input epochs of the 28 participants included in the EEG analyses) underlying the reported analyses, together with the stimulus lists and experiment programs, are publicly available in an OSF repository at https://doi.org/10.17605/OSF.IO/TSA67 (Liang, 2026). Raw EEG recordings are not publicly deposited because they contain identifying information.

## Code availability

Custom analysis code for EEG preprocessing, NPS searchlight computation, TFCE permutation inference, the multilevel mediation models (R and Mplus; Mplus requires a separate licence), the written-response revision models and the model-based representational similarity analysis is available in the same OSF repository under the MIT licence. Software versions are listed in Supplementary Methods SM11.

## Acknowledgements

We thank all participants. We thank Y. Hu (Chongqing University of Arts and Sciences) for organising student helpers for data collection, X. Liu (Chongqing University of Arts and Sciences) for arranging accommodation and meals during data collection, and X. Wu and W. Wang (Chongqing University of Arts and Sciences) for help with data collection. Large language model assistants (Claude, Anthropic; ChatGPT and Codex, OpenAI) were used to help write and check code, screen the literature and edit the text, as described in Methods.

## Author contributions

Y.L. conceived and designed the study, collected and analysed the data, prepared the figures and wrote the manuscript. Y.H. contributed to the early experimental design and the selection of materials. Z.C. programmed the early version of the experiment. Y.Li contributed to writing and discussion of the manuscript in its later stage. L.L. helped with the literature search and discussed the statistical methods. Y.W., Y.S., F.R. and M.S. collected the data. W.L. contributed to the early discussion of the experimental design, funded part of the participant payments and coordinated data collection at Chongqing University of Arts and Sciences. X.Y. contributed to revising the manuscript. L.Z. provided guidance on the statistical approach. All authors approved the final manuscript.

## Ethics declarations

### Human ethics approval

This study involved human participants and was approved by the Institutional Review Board of Chongqing University of Arts and Sciences, where the data were collected. It was conducted in accordance with the Declaration of Helsinki.

### Consent to participate

Written informed consent to participate was obtained from all participants after the EEG procedures and the voluntary nature of participation had been explained.

