## Supplementary Information for "Neural mechanisms by which spaced learning promotes understanding"

---

#### Supplementary Methods

##### SM1. Participant exclusion criteria

Epochs with residual artefacts were repaired or rejected with AutoReject (SM6). Participant-level EEG quality was judged by visual inspection of each participant's evoked responses; four participants whose evoked waveforms showed a low signal-to-noise ratio or an unstable pre-stimulus baseline were excluded. No fixed epoch-rejection threshold was applied at the participant level.

Six participants were excluded on the basis of laboratory records made during the sessions: one whose final session took place a day later than scheduled; one who reported changing memorisation strategy mid-session; one whose final review was hurried; two whose sessions had excessive ambient noise, one of which was also unusually short; and one whose session was repeatedly interrupted and whose initial-test records were incomplete.

Six further participants were not analysed: five who had run an earlier version of the test program, and one whose session was interrupted by a power failure, after which they reported marked emotional distress.

---

##### SM2. Stimuli selection and validation

English words were drawn from GRE vocabulary lists (Yu, 2010), excluding CET-4 items (New Oriental Test Research Centre, 2016). One rater who had passed CET-6 screened the remaining pool and selected 200 words they reported as unfamiliar, with etymological explanations confirmed as reasonable by two additional CET-6 students. These 200 words were assessed for familiarity by 27 undergraduates who did not participate in the main experiment (CET-4 scores: 430–550).

**Assessment procedure.** In each trial, participants first viewed an English word and judged whether they knew its meaning (yes/no). To distinguish transient recall failure from true unfamiliarity, the word was then presented together with its Chinese translation, and participants rated their familiarity on a 1–5 scale (1 = completely unfamiliar; 5 = highly familiar).

**Selection and grouping.** Familiarity ratings were summed across all 27 raters and sorted. The 150 words with the lowest total familiarity scores were selected for the main experiment and divided into three sets using equal-interval allocation. Two alternative partitions of the 150 words into three sets were used across the sample. For each participant, one set served as the once-learning (O) condition and the other two were assigned to the spaced (S) and massed (M) conditions, with this assignment counterbalanced across participants. The three word sets did not differ significantly in summed familiarity, $F(2,147) < 1$, or word length, $F(2,147) = 1.38$, $P = 0.25$. To reduce learning difficulty, the 50 words assigned to each condition were divided into two sets of 25 words. An additional 100 words, drawn at random from the same GRE lists, served as novel foils for the old–new recognition test.

**Stimulus structure.** Each word pair included the English word, its Chinese translation, a structural analysis decomposing the word into familiar simple cognates (confirmed to belong to CET-4 vocabulary), and an etymological derivation linking the cognates to the meaning. Representative examples are shown in Supplementary Table 9.

#### SM3. Mastery-gated learning and design alternatives

Learning was mastery-gated (Methods): study time was free to vary while criterion-verified completion was held constant across conditions. We prioritised this criterion-verified endpoint over equal study duration because the two controls could not be imposed simultaneously. A fixed duration calibrated to the shorter massed-learning requirement could stop spaced learning before criterion, whereas a duration calibrated to the longer spaced-learning requirement would force continued massed study after criterion had already been reached. A yoked-duration design would create the same problem at the participant level. Both alternatives would therefore compare conditions at different learning endpoints or impose post-criterion overlearning, reducing ecological validity. Self-paced, criterion-gated learning instead mirrors the way learners ordinarily continue until they can produce the material, while allowing the additional effort associated with delayed restudy to be observed (Bjork & Bjork, 1992; Kornell & Bjork, 2008). The resulting duration difference is a measured cost of reaching a common, criterion-verified endpoint, not by itself evidence for a particular mechanism.

Changing from a within- to a between-participant design would address only part of the remaining temporal structure. A two-group spaced-versus-massed design would remove direct cross-condition carryover within a participant, but the groups would still differ in the practice and strategy-development opportunities created by their schedules: the same two encounters would be separated across days for spacing and compressed within one session for massing. Adding a third once-learning group would provide the one-scheduled-episode reference required for baseline-relative NPS, but it would not separate these schedule-contingent changes from spacing, because one, massed, and spaced learning necessarily entail different encounter histories. Extensive pretraining until strategy use reached a common asymptote could further reduce this variation, but it would change the target question to spacing after task-specific training rather than spacing during ordinary mastery-gated learning.

#### SM4. Behavioural data analysis: statistical specifications

For the within-subject paired comparisons reported in the main text we report Cohen's $d_z$, computed as the mean difference divided by the standard deviation of the difference scores (using an $n-1$ denominator), with 0.2, 0.5, and 0.8 denoting small, medium, and large effects. We use $d_z$ because it accounts for the correlation between paired observations.

To control the family-wise error rate across the three pairwise comparisons of each outcome (S vs M, S vs O, M vs O), we applied the Holm-Bonferroni sequential correction, ordering p-values from smallest to largest and testing each against its sequential $\alpha$ threshold at a family-wise $\alpha$ of 0.05.

---

#### SM5. Rationale and inferential boundary of final-test EEG

EEG was recorded during the Day 5 final test, not during the learning phases (scheduled Days 1 and 3). This design placed all three schedules in the same recording context after the common Day 3-to-Day 5 interval. It therefore addressed how schedule-linked organisation was expressed during delayed retrieval; it could not identify when that organisation formed.

A second encounter in repeated learning necessarily combines re-exposure with retrieval of the first encounter. Recording only the final test avoided comparing EEG across learning sessions and supplied a common observation point. The once-learning condition additionally supplied the one-scheduled-episode reference used for baseline-relative NPS. Neither feature isolates a pure retrieval process: final retrieval reflects the accumulated learning history, and the study contains no encoding-stage neural measurement.

Within this boundary, the corrected late NPS field identifies schedule-linked differentiation observable at retrieval, and the NN indirect effect identifies a partly overlapping set of cells statistically associated with the relative delayed advantage in self-rated explanatory retrieval fluency. It does not show that spacing's benefit was created at retrieval. Future studies recording both learning and retrieval could test how encoding-stage changes are carried into final retrieval.

---

#### SM6. EEG preprocessing parameters

Before ICA, the continuous data were band-pass filtered at 0.1–30 Hz with a zero-phase finite-impulse-response (FIR) filter (MNE default design; 0.1-Hz transition band at the high-pass edge, filter order determined automatically) and notch-filtered at 50, 100, 150 and 200 Hz.

Independent component analysis (ICA) was performed in MNE-Python (`mne.preprocessing.ICA`) on a copy of the data high-pass filtered at 1 Hz, decomposing the signal into 50 components (fixed random seed). Ocular components were identified automatically with `find_bads_eog` (z-score threshold = 3.0) using the

VEOG channel, and the flagged components were confirmed by visual inspection of their topographies and source time courses before being removed from that same 1-Hz high-passed copy, which was retained as the analysed signal.

Data were recorded and analysed at 500 Hz; no resampling was applied.

After epoching, residual artefacts were handled with AutoReject (Jas et al., 2017). Channel-wise peak-to-peak thresholds were learned by Bayesian optimisation, and cross-validation selected the maximum number of channels to interpolate within an epoch (candidates 1–15) and the consensus fraction of bad channels above which an epoch was rejected (ten candidates from 0 to 0.25); the random seed was fixed (23).

#### SM7. Trial selection and balancing

Trials entered the NPS analysis only when they met all of the following criteria: a correct old/new judgement (recognition = 1 for studied items), a confidence rating ≥ 3 on the 0–5 scale, and an epoch retained after artefact handling (SM6). Completeness of the first-session covariates was not a selection criterion; one selected massed trial lacked a first-session logical score and was therefore excluded from the mediation models (SM9).

To keep trial counts comparable across conditions within each participant, the number of once-learning trials surviving the inclusion criteria defined the participant's target count. For the massed and spaced conditions, trials were ranked by final logical score and the highest-scoring trials were retained up to that target; a condition with fewer eligible trials kept all of them. Counts were therefore equal across the three conditions in 24 of the 28 participants and differed by 1–10 trials in the remaining four (x009: O 34, M 31, S 34; x037: 29, 29, 27; x038: 44, 34, 39; x047: 38, 37, 38). In total, 932 once-learning, 918 massed and 925 spaced trials were retained (mean 33.0 per participant and condition; range 22–46). In the massed condition the two learning episodes were separated by a median of 2.2 h (interquartile range 1.7–2.7 h; measured from the learning-session files), during which participants completed the once-learning and spaced-restudy sets; a few participants left the laboratory briefly during the Day 3 session, so the longest intervals reflect breaks rather than continuous study.

#### SM8. NPS computation: parameter justification

The spatial searchlight used a 3D Euclidean radius of 5 cm, computed from the three-dimensional electrode coordinates, which balances spatial specificity against sufficient channel coverage; typical searchlight patches contained 5–8 electrodes. Smaller radii (e.g., 2 cm) risk including too few channels for stable multivariate patterns, whereas larger radii (e.g., 10 cm) sacrifice spatial resolution. Four temporal-pole electrodes (FT9, FT10, TP9, TP10) were excluded because of their high susceptibility to muscle artefacts and poor signal-to-noise ratio, following standard practice in high-density EEG studies.

Patterns were extracted in sliding temporal windows 100 ms wide, stepped in 10 ms increments, giving 91 windows spanning the −200 to 800 ms epoch. The 100 ms width captures transient ERP components while retaining adequate temporal resolution, consistent with prior EEG decoding work (King & Dehaene, 2014), and the 90% overlap between successive windows yields smooth temporal dynamics at acceptable computational cost.

Neural pattern similarity was quantified as the Pearson correlation between condition patterns, with higher values indicating greater similarity. We preferred Pearson r over Euclidean distance because it is scale-invariant and robust to amplitude differences, over cosine similarity because it mean-centres the patterns and is thus more sensitive to pattern differences, and over Mahalanobis distance because it requires no covariance-matrix estimation.

---

#### SM9. Multilevel mediation: model specification

The two-level mediation model (Methods) comprised two regression equations and one derived quantity:

$NPS_c = \beta_0 + a \cdot \mathrm{Condition} + \beta_1 \cdot C_1 + \beta_2 \cdot C_2 + u + \varepsilon_1$ (1)

$\mathrm{LogicAdv} = \gamma_0 + b \cdot NPS_c + c' \cdot \mathrm{Condition} + \gamma_1 \cdot C_1 + \gamma_2 \cdot C_2 + v + \varepsilon_2$ (2)

Indirect effect = $a \times b$ (3)

In equation (1), the grand-mean-centred mediator $NPS_c$ was regressed on condition, yielding the *a*-path. In equation (2), LogicAdv—the target item's final-test logical advantage relative to the once-learning baseline, computed for each trial pair as the target-word logical score minus the paired baseline-word logical score—was regressed on $NPS_c$ (the *b*-path) and condition (the direct effect, $c'$), adjusting for the same covariates. Here $\beta_0$ and $\gamma_0$ are intercepts; $\beta_1$, $\beta_2$, $\gamma_1$, and $\gamma_2$ are covariate coefficients; $C_1$ and $C_2$ are the grand-mean-centred first-session logical score and study-duration covariates; *u* and *v* are participant-level random intercepts; and $\varepsilon_1$ and $\varepsilon_2$ are residual terms.

---

#### SM10. TFCE permutation inference: implementation details

Spatial adjacency was defined by Delaunay triangulation of the 2D electrode coordinates (implemented with `scipy.spatial.Delaunay`), producing a graph connecting neighbouring electrodes; for formal inference this graph was restricted to the 18 centres of the bilateral central domain, which left no adjacency between hemispheres. Temporal adjacency connected successive time windows, which overlap given the 10 ms step and leave no temporal gaps. Combining the two, each spatiotemporal unit (electrode *i*, time *t*) was connected to its spatially adjacent electrodes at the same time *t* and to the same electrode at the adjacent times $t \pm 1$.

The two path families were corrected separately, and the 95th percentile of a null distribution was used only as a descriptive summary rather than as the decision rule. The completed permutation archive contained 1,992 maps: the observed map (permutation 0) and permutations 1–1,991 (1–907 from the original run and 908–1,991 from an extension run under the same shuffling procedure). Audit invariants were constant across all maps: 28 participants, 1,843 participant-by-target blocks, and 63,294 input trial-pair rows; participant-level condition counts were preserved and no target block was split. Because the exchangeability operation randomised the condition labels, this procedure calibrated the condition-label null branch; although $b$ was re-estimated in every refit, mediator-outcome pairing was not independently randomised to construct a separate $b$-null. The number of permutations was limited by computation: each permutation required re-fitting the full MLR mediation model at all 1,278 units (18 centres × 71 window centres), so it was set by the available computing time rather than by the results. Monte Carlo uncertainty did not affect the peak-level results: the exact binomial 95% confidence intervals of the minimum corrected $P$ values lay entirely below 0.05 (path $a$, 0.016–0.029; NN, 0.016–0.030).

---

#### SM11. Software and code availability

EEG preprocessing and ICA were run in MNE-Python 0.23 (FastICA, 50 components, max_iter = 200). Subsequent analyses were performed in Python 3.10.11 (MNE-Python 1.8.0, NumPy 1.23.5, SciPy 1.10.1, pandas 2.3.3, statsmodels 0.14.6), which handled epoching and AutoReject, NPS computation, TFCE permutation inference, and the behavioural mixed-effects models (LMM/GEE). The multilevel mediation models were fit in Mplus 8.3 (Muthén & Muthén, 1998–2017), called from R 4.4.2 via the MplusAutomation package, with parallelisation implemented with the base parallel package (makeCluster/parLapplyLB). Figures were produced with matplotlib 3.10.8; seaborn 0.12.2 was installed in the same environment and used for exploratory plots only.

Access to data, materials and custom analysis code is described in the Data availability and Code availability sections of the main article.

---

#### SM12. Representational similarity analysis: schedule contrasts

Direct contrasts between schedules were run for CORnet-Z over the 17 posterior patches. For each participant and each of the three contrasts—spaced minus once-learning (tested negative), massed minus once-learning and massed minus spaced (both tested positive)—the paired difference in Fisher-$z$ RSA was regressed on the matching differences in first-session logical score and first-session study duration (scaled, not centred), and the adjusted intercept $t$ (residual df = 25) formed the statistic at every one of the 1,547 cells. Threshold-free cluster enhancement ($H = 2$, $E = 1$, 1,000 integration steps) used the same spatial and temporal adjacency as the main analyses. The null was built from 5,000 whole-participant sign flips of the reduced-model residuals (Freedman–Lane), giving $p = [1 + \text{count}(\text{null maxima} \geq \text{observed})]/5{,}001$. Each contrast × stage combination was corrected as its own family ($3 \times 4 = 12$ families), with no correction across families. Because the sign-flip null is conditional on the fitted nuisance model, the resulting family-wise control is approximate rather than exact.

---

### Supplementary Results

#### S1. Data exploration and distribution

The sample comprised 28 participants contributing 4,199 trials (spaced, 1,399; massed, 1,400; once-learning, 1,400; one spaced trial of one participant was not recorded), with CET-4 scores ranging from 400 to 542. Recognition in the spaced condition showed a ceiling tendency (~97.3%), the study-duration measures (first- and second-session) were right-skewed with dispersion varying across conditions, and logical scores were more symmetrically distributed.

Shapiro-Wilk statistics are listed in Supplementary Table 2; the combination of paired comparisons, multilevel models and GEE (for binary recognition) accommodates the observed departures from normality.

---

#### S2. First-session condition comparisons

For first-session logical score, the Friedman omnibus test was non-significant ($\chi^2$=2.62, $P = 0.270$), and no pairwise comparison survived correction (S vs M: t(27)=2.05, $P = 0.050$, $P_{\text{Holm}} = 0.151$, $d_z$=0.39; S vs O: t(27)=1.23, $P = 0.231$, $P_{\text{Holm}} = 0.461$, $d_z$=0.23; M vs O: t(27)=-0.16, $P = 0.874$, $P_{\text{Holm}} = 0.874$, $d_z$=-0.03). Thus, the measured first-session logical score did not differ detectably across conditions.

For first-session study duration, the omnibus test was significant ($\chi^2$=27.64, $P < 0.001$), with all pairwise comparisons differing (S vs M: t(27)=3.66, $P = 0.001$, $P_{\text{Holm}} = 0.002$, $d_z$=0.69; S vs O: t(27)=7.25, $P < 0.001$, $P_{\text{Holm}} < 0.001$, $d_z$=1.37; M vs O: t(27)=3.06, $P = 0.005$, $P_{\text{Holm}} = 0.005$, $d_z$=0.58). These differences reflect practice effects inherent to the within-subject learning sequence (S items learned first, O items last) rather than a randomisation failure; the trial-level models adjust for them (Supplementary Results S4).

---

#### S3. Robustness checks

Shapiro-Wilk tests were conducted separately by condition to evaluate the distributional assumptions behind the paired parametric summaries (Supplementary Table 2). Logical scores were generally approximately normal, whereas recognition deviated from normality in a ceiling-related way in the spaced and massed conditions. Because recognition accuracy approached ceiling in the spaced condition (~97%) and was non-normal in both spaced and massed conditions, we supplemented the parametric recognition comparisons with Wilcoxon signed-rank tests (S vs M: W=47.0, $P = 0.010$; S vs O: W=0.0, $P < 0.001$; M vs O: W=0.0, $P < 0.001$), each consistent with its parametric counterpart. The recognition results were therefore robust to distributional assumptions. We limited non-parametric follow-up to recognition, the outcome whose non-normality was ceiling-related; for final logical score and confidence, inference rests on the paired parametric summaries and the covariate-adjusted trial-level LMM/GEE models reported in Supplementary Table 4.

---

#### S4. Complete trial-level model results (LMM/GEE)

Trial-level models were fitted in three steps (O = reference unless noted). For final logical score and recognition, Step 1 included condition only; from Step 1 onward, the confidence models additionally included recognition and the condition × recognition interaction. Step 2 added the grand-mean-centred first-session study duration and first-session logical score as covariates to all three outcome models, and Step 3 added the condition × CET-4 (centred) moderation term. Final logical score and confidence were modelled with linear mixed-effects models (LMMs) including participant-specific random intercepts. Recognition was modelled with generalised estimating equations (GEE) using a binomial distribution and logit link, participant as the clustering variable, and an exchangeable working correlation structure. The condition contrasts reported in the main text and Supplementary Table 4 are the Step-3 condition coefficients evaluated at mean CET-4; for confidence, the condition coefficients are read with recognition grand-mean centred (the overall adjusted contrast), and a second model restricted to correctly recognised trials, without the recognition term, gives the contrast among recognised words; confidence was not thresholded in that model so as not to truncate the outcome. CET-4 moderation terms are reported in Supplementary Table 5.

For final logical score and recognition, the large spaced and massed advantages over the once-learning baseline were essentially unchanged when first-session study duration and first-session logical score were added as covariates. For final logical score (LMM), the spaced-versus-once advantage shifted only from b=2.53 (Step 1) to b=2.68 (Step 2; z=24.33, $P < 0.001$) and the massed-versus-once advantage from b=1.85 to b=1.92 (Step 2; z=17.81, $P < 0.001$); for recognition (GEE), the corresponding values were b=2.08 → 2.17 ($P < 0.001$) and b=1.35 → 1.38 (Step 2; $P < 0.001$). The covariate-adjusted Step-3 spaced-versus-massed contrast, evaluated at mean CET-4, remained significant for final logical score (b=0.759, z=6.99, $P < 0.001$) and recognition (GEE b=0.718, $P < 0.001$), whereas for confidence the overall adjusted contrast (recognition centred) was not significant (b=0.058, z=1.65, $P = 0.098$) and the contrast among correctly recognised words was (b=0.083, z=2.77, $P = 0.006$). Full coefficients are summarised in Supplementary Table 4.

**Descriptive paired comparisons.** Unadjusted spaced-minus-massed comparisons of participant-level summaries yielded Cohen's $d_z$ values of 0.83 for final logical score, and 0.56 for studied-item hit rate (N = 28; see Supplementary Methods SM4 for the paired-effect definition). For final logical score, unadjusted paired comparisons with once-learning yielded $d_z = 2.47$ for spaced learning and 2.62 for massed learning. These values describe the unadjusted paired comparisons rather than standardised versions of the covariate-adjusted model estimates.

#### S5. Methodological transparency

A sensitivity calculation for a two-tailed paired t-test ($\alpha = 0.05$, power = 0.80) shows that N = 24 detects a medium within-participant effect ($d_z = 0.60$, based on the Cepeda et al., 2006 meta-analysis); the achieved sample of N = 28 detects effects of $d_z \geq 0.55$ at the same power.

The study was not pre-registered. Planned analyses comprised Friedman tests, paired t-tests, and effect sizes; we added post hoc the LMM/GEE models with baseline covariates (after observing the first-session study-duration differences), the CET-4 moderation analysis (exploratory but theoretically motivated), and the non-parametric robustness checks (after observing ceiling effects). For multiple comparisons, the three pairwise comparisons per outcome (S vs M, S vs O, M vs O) were Holm-Bonferroni corrected; p values for the trial-level model coefficients were not corrected for multiplicity. Confidence interaction analyses are explicitly labelled as exploratory and should be interpreted with caution. The two confidence readings (recognition centred; correctly recognised trials only) were specified after the primary analyses, prompted by the recognition-based selection of EEG trials.

#### S6. Representational similarity analysis: schedule contrasts

Supplementary Table 10 summarises the temporal support of the single-condition CORnet-Z tests shown in Figure 5.

None of the twelve CORnet-Z contrast families returned a corrected cell. Across the three contrasts and four stages, the smallest family-wise corrected $p$ was .194 (spaced minus once-learning at V4); the largest was .922. The stage-wise asymmetry visible in the single-condition maps—early-stage support present under once-learning and massed but not under spaced—is therefore a description of where each condition's RSA exceeded zero, not a tested difference between conditions.

#### S7. Written-response revision: rate, size and convergence

At each initial test, participants wrote the intermediary words and the Chinese definition for every target. For the two repeated conditions, a word was scored as revised when the normalised written response (Unicode NFKC, whitespace collapsed; no case folding) at the second initial test differed from that at the first; the comparison records whether the response changed, not whether it improved. Of the 760 revisions, 14 differed only in letter case, punctuation, cue order, or an accompanying Chinese gloss (12 spaced, 2 massed); they are retained, because the measure is defined on what was written.

A word contributed to the measures only when both responses were observed (2,550 words). Two of the 28 participants fell below 20 such words in at least one condition (one with 0 observable words under massed and 3 under spaced; the other with 1 under massed and 0 under spaced) and are therefore absent from these measures, leaving 26. The threshold was applied to written-record availability before and independently of outcome assessment.

**Primary inference (word-level mixed models; Methods; pipeline in Supplementary Figure 2).** Revision was more frequent after spaced than massed learning (35.2% versus 24.9% of observable words, participant means; pooled proportions 35.1% and 24.4%): adjusted odds ratio 1.75 (95% CI [1.44, 2.13], $z = 5.57$, $P = 2.5 \times 10^{-8}$; random-intercept variances: participants 0.74, words 0.63). Among revised words, revisions were smaller after spaced learning: adjusted spaced-minus-massed difference in embedding distance −0.021 ([−0.037, −0.005], $t(716) = -2.62$, $P = 0.009$; variances: participants 0.0007, words 0.0015, residual 0.0105). Revised responses converged on other learners' formulations: the T1-to-T2 change in distance to the same-test group centre differed by −0.027 ([−0.047, −0.007], $P = 0.007$; unadjusted means: spaced −0.010, massed +0.012). At T1 the two schedules did not differ in distance to the group centre (−0.007 [−0.019, +0.006], $P = 0.28$); at T2 they did (−0.033 [−0.048, −0.019], $P < 0.0001$). For the convergence outcomes the word-intercept variance was estimated at zero, so the crossed and participant-only models coincide.

**Sensitivity analyses specified with the primary models.** With participant-only random intercepts: odds ratio 1.58 ([1.31, 1.90], $P = 1.4 \times 10^{-6}$); size −0.023 ([−0.040, −0.007], $P = 0.006$). Under the alternative covariate convention (first-session logical score and log study duration centred within participant and condition, no CET-4 term): odds ratio 1.86 ([1.54, 2.26], $P = 2.6 \times 10^{-10}$; participant-only 1.78 [1.48, 2.14]); size −0.019 ([−0.034, −0.003], $P = 0.019$; participant-only −0.021 [−0.037, −0.004], $P = 0.013$). All eight models converged without singular fits. Revision size computed from four alternative embedding models on the intermediary-word field alone (565 revised words with that field) gave the same sign in every case (MiniLM-L12 −0.030, $P = 0.11$; BGE-base-zh −0.015, $P = 0.30$; BGE-M3 −0.018, $P = 0.11$; Qwen3-Embedding-8B −0.017, $P = 0.045$), all four less precise than the primary model; effect sizes are not compared across embedding models because they operate on different distance scales.

**Convergence: specification and robustness.** Against a fixed centre (both tests measured against the other learners' T2 centre), the contrast was −0.028 ([−0.047, −0.008], $P = 0.007$). The contrast was insensitive to the reference: other learners' T1 centre −0.026 ([−0.046, −0.007], $P = 0.008$); pooled T1 and T2 centre −0.027 ($P = 0.007$); centre built only from learners who studied the word under the other schedule −0.024 ([−0.044, −0.004], $P = 0.020$, 748 words); only from the same schedule −0.029 ($P = 0.009$). Adding a by-participant random slope for condition gave −0.029 ($P = 0.012$). A within-participant permutation test (10,000 shuffles of the schedule labels within participant; word sets were counterbalanced across participants, Methods) gave $P = 0.019$ at the word level and $P = 0.015$ for the participant-level mean difference. In 100 random split-halves of the words, the rate effect kept its sign and $P < 0.05$ in all 200 halves (5th–95th percentile of the odds ratio 1.49–2.07), the size effect kept its sign in all 200 halves ($P < 0.05$ in 56%), and the convergence effect kept its sign in 199 of 200 halves ($P < 0.05$ in 48%).

The measures index observable change in what participants wrote and where it moved relative to other learners' responses. They do not establish improved explanations, semantic reorganisation, or mediation of later outcomes.

---

### Supplementary Figures

#### Supplementary Figure 1. Unsubtracted NPS time courses in the bilateral central domain

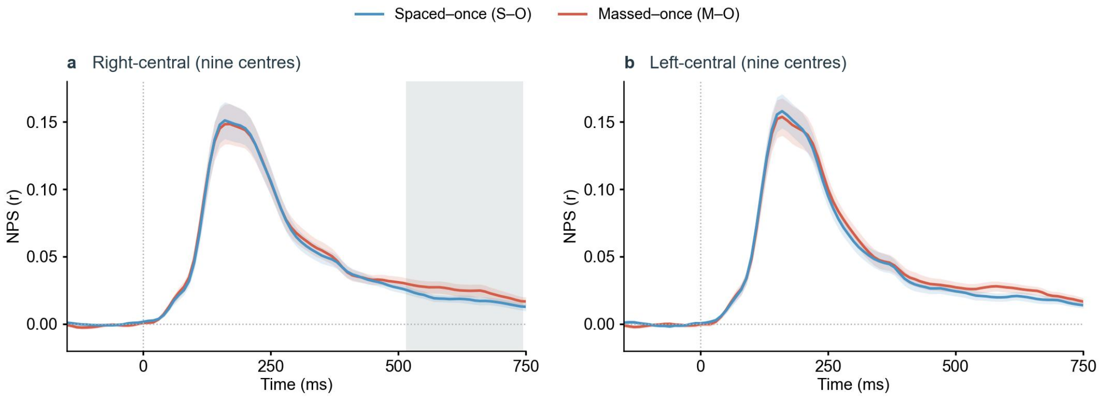


*Supplementary Figure 1. Unsubtracted NPS time courses in the bilateral central domain.* For each participant, the trial-mean NPS between final-test trials and once-learning trials for spaced (S–O, blue) and massed (M–O, red) items, averaged over the nine right-central (**a**) and nine left-central (**b**) searchlight centres of the bilateral central domain; lines and bands, mean ± s.e.m. across participants (N = 28); 91 window centres from −150 to 750 ms (100-ms windows stepped by 10 ms); 0, word onset. Both time courses are positive at every window centre from 50 to 750 ms on both sides. The grey band in **a** marks the window centres with corrected path-a support in Figure 4a (520–740 ms).

#### Supplementary Figure 2. Written-response revision pipeline: comparison and convergence

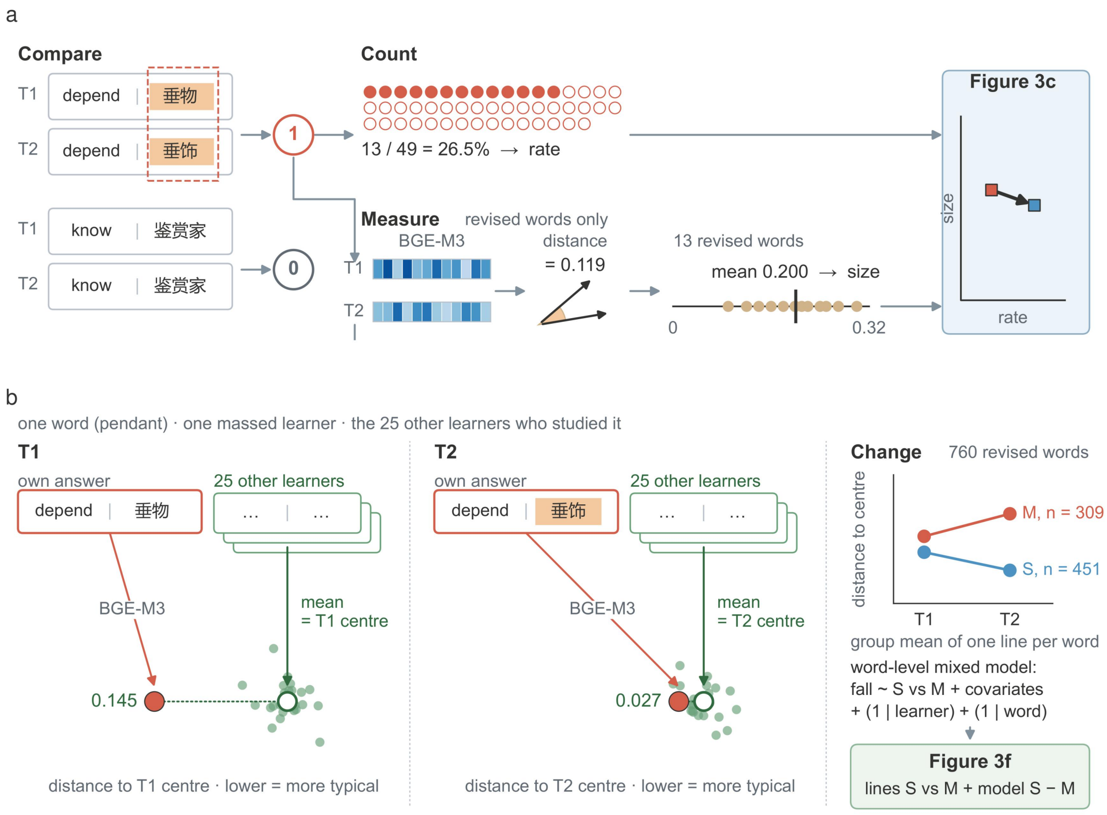


*Supplementary Figure 2. Written-response revision pipeline: comparison and convergence.* **a**, Comparison within a learner. For each word, the T1 and T2 written responses (intermediary words | Chinese definition) are compared after normalisation; a word counts as revised (1) when the joint text changed and as unrevised (0) otherwise. Counting over a learner's observable words gives the revision rate (example learner: 13 of 49 words). For revised words only, both responses are embedded (BGE-M3) and the cosine distance between the T1 and T2 embeddings gives the revision size; the learner's mean over revised words is the size shown in Figure 3c. **b**, Convergence against other learners, illustrated for one word (pendant) and one massed learner. At each test, the other learners' responses to the same word are embedded and averaged (leave-one-out) to give that test's group centre; the learner's own response is placed in the same space and its distance to the centre is measured (T1 against the T1 centre, T2 against the T2 centre; lower = more typical of the group). The change in this distance from T1 to T2 is the convergence outcome; Figure 3f shows the group means of one such line per revised word (spaced, n = 451 words; massed, n = 309) and the word-level mixed model tests the spaced-minus-massed difference in the change. Example values are from the frozen result tables; the point clouds are schematic.

#### Supplementary Figure 3. Delayed recognition confidence among correctly recognised words

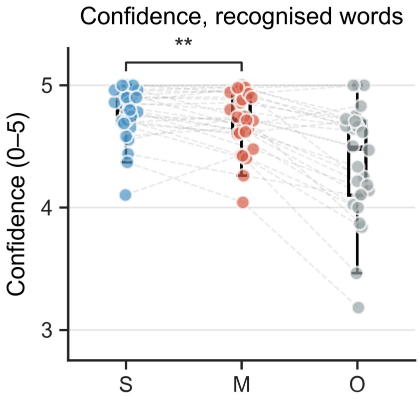


*Supplementary Figure 3. Delayed recognition confidence among correctly recognised words.* Participant means of the 0–5 confidence rating over correctly recognised studied words at the Day-5 final test, for spaced (S), massed (M) and once-learning (O) items (N = 28; dots, participants; grey lines, within-participant pairs; box, median and interquartile range). Bracket: spaced-versus-massed contrast from the LMM fitted to correctly recognised trials ($b$ = 0.083, $P$ = 0.006; Supplementary Table 4). Over all trials, with recognition centred, the contrast was $b$ = 0.058, $P$ = 0.10.

---

### Supplementary Tables

#### Supplementary Table 1. First-Session Condition Comparisons

| Variable | Friedman $\chi^2(2)$ | p | S vs M | S vs O | M vs O |
|---|---|---|---|---|---|
| Logical score (T1) | 2.62 | 0.270 | $P$ = 0.151 | $P$ = 0.461 | $P$ = 0.874 |
| Study duration (T1) | 27.64 | <0.001 | $P$ = 0.002 | $P$ < 0.001 | $P$ = 0.005 |
| Production accuracy (T1) | N/A | N/A | N/A | N/A | N/A |

*Note*: All pairwise comparisons use Holm correction. Production accuracy (T1) not tested (100% in all conditions by the stopping rule); no recognition test was given at T1.

---

#### Supplementary Table 2. Normality Tests (Shapiro-Wilk)

| Variable | Condition | Shapiro-Wilk W | p | Interpretation |
|---|---|---|---|---|
| Logical score (T1) | S | 0.965 | 0.447 | Approximately normal |
| Logical score (T1) | M | 0.984 | 0.936 | Approximately normal |
| Logical score (T1) | O | 0.972 | 0.631 | Approximately normal |

| Variable | Condition | Shapiro-Wilk W | p | Interpretation |
|---|---|---|---|---|
| Logical score (T2) | S | 0.961 | 0.360 | Approximately normal |
| Logical score (T2) | M | 0.978 | 0.791 | Approximately normal |
| Final logical score | S | 0.923 | 0.042 | Mild deviation |
| Study duration (T1) | M | 0.895 | 0.009 | Deviation |
| Study duration (T1) | O | 0.837 | <0.001 | Deviation |
| Study duration (T2) | S | 0.924 | 0.044 | Mild deviation |
| Confidence | S | 0.878 | 0.004 | Deviation |
| Recognition | S | 0.893 | 0.008 | Ceiling-related deviation |
| Recognition | M | 0.884 | 0.005 | Ceiling-related deviation |

---

#### Supplementary Table 3. Recognition Non-Parametric Robustness Checks

| Comparison | Outcome | Wilcoxon W | p | Parametric conclusion | Agreement |
|---|---|---|---|---|---|
| S vs M | Recognition | 47.0 | 0.010 | Significant S > M | Yes |
| S vs O | Recognition | 0.0 | <0.001 | Significant S > O | Yes |
| M vs O | Recognition | 0.0 | <0.001 | Significant M > O | Yes |

*Note*: Non-parametric follow-up tests were restricted to recognition because its deviation from normality was ceiling-related. Final logical score and confidence are reported using paired parametric summaries and covariate-adjusted trial-level models.

---

#### Supplementary Table 4. Trial-Level Model Fixed Effects (Step 3)

All models adjusted for first-session study duration and first-session logical score (both grand-mean-centred). Final logical score and confidence: LMM with subject random intercepts. Recognition: GEE, binomial-logit. Confidence models additionally included recognition and condition × recognition, with recognition grand-mean centred so that the condition coefficients give the overall adjusted contrast; a further block gives the model fitted to correctly recognised trials only (no recognition term). Each block lists every fixed effect of the Step-3 model with once-learning (O) as the reference and CET-4 grand-mean-centred, so condition coefficients are evaluated at mean CET-4; these are the values reported in the main text. The final row of each block gives the spaced-versus-massed contrast from the same model refitted with massed (M) as the reference. b, unstandardised coefficient (log-odds for recognition); SE, standard error; z, Wald statistic; CI, confidence interval.

**Final logical score (LMM)**

| Term | b | SE | z | *P* | 95% CI |
|---|---|---|---|---|---|
| Intercept | 5.346 | 0.197 | 27.09 | < 0.001 | [4.959, 5.733] |
| Massed vs Once | 1.918 | 0.108 | 17.82 | < 0.001 | [1.707, 2.129] |
| Spaced vs Once | 2.677 | 0.110 | 24.33 | < 0.001 | [2.462, 2.893] |
| CET-4 (centred) | -0.013 | 0.006 | -2.07 | = 0.038 | [-0.025, -0.001] |
| Massed vs Once × CET-4 (centred) | 0.005 | 0.003 | 1.55 | = 0.121 | [-0.001, 0.012] |
| Spaced vs Once × CET-4 (centred) | 0.002 | 0.003 | 0.66 | = 0.509 | [-0.004, 0.009] |
| First-session study duration (centred) | -0.014 | 0.002 | -8.12 | < 0.001 | [-0.018, -0.011] |
| First-session logical score (centred) | 0.166 | 0.025 | 6.70 | < 0.001 | [0.117, 0.214] |
| Spaced vs Massed (same model, Massed as reference) | 0.759 | 0.109 | 6.99 | < 0.001 | [0.546, 0.972] |

**Recognition (GEE, binomial-logit)**

| Term | b | SE | z | *P* | 95% CI |
|---|---|---|---|---|---|
| Intercept | 1.460 | 0.146 | 10.00 | < 0.001 | [1.174, 1.746] |
| Massed vs Once | 1.443 | 0.119 | 12.16 | < 0.001 | [1.210, 1.676] |
| Spaced vs Once | 2.161 | 0.167 | 12.97 | < 0.001 | [1.834, 2.487] |
| CET-4 (centred) | 0.000 | 0.004 | -0.10 | = 0.920 | [-0.008, 0.008] |
| Massed vs Once × CET-4 (centred) | 0.013 | 0.004 | 3.18 | = 0.001 | [0.005, 0.021] |
| Spaced vs Once × CET-4 (centred) | -0.002 | 0.004 | -0.51 | = 0.608 | [-0.011, 0.006] |
| First-session study duration (centred) | -0.006 | 0.002 | -3.29 | < 0.001 | [-0.010, -0.003] |
| First-session logical score (centred) | 0.002 | 0.031 | 0.06 | = 0.954 | [-0.059, 0.062] |
| Spaced vs Massed (same model, Massed as reference) | 0.718 | 0.178 | 4.04 | < 0.001 | [0.369, 1.066] |

**Confidence (LMM; recognition grand-mean centred — overall adjusted contrast)**

| Term | b | SE | z | *P* | 95% CI |
|---|---|---|---|---|---|
| Intercept | 4.225 | 0.062 | 67.86 | < 0.001 | [4.103, 4.347] |
| Massed vs Once | 0.281 | 0.034 | 8.22 | < 0.001 | [0.214, 0.348] |
| Spaced vs Once | 0.339 | 0.036 | 9.44 | < 0.001 | [0.269, 0.409] |
| Recognition (centred) | 1.595 | 0.062 | 25.78 | < 0.001 | [1.474, 1.717] |

| Term | b | SE | z | *P* | 95% CI |
|---|---|---|---|---|---|
| Massed vs Once × Recognition (centred) | 0.591 | 0.121 | 4.90 | < 0.001 | [0.355, 0.828] |
| Spaced vs Once × Recognition (centred) | 0.815 | 0.158 | 5.16 | < 0.001 | [0.506, 1.125] |
| CET-4 (centred) | -0.002 | 0.002 | -0.91 | = 0.363 | [-0.006, 0.002] |
| Massed vs Once × CET-4 (centred) | 0.002 | 0.001 | 1.93 | = 0.054 | [0.000, 0.004] |
| Spaced vs Once × CET-4 (centred) | 0.002 | 0.001 | 1.49 | = 0.135 | [0.000, 0.004] |
| First-session study duration (centred) | -0.002 | 0.001 | -4.16 | < 0.001 | [-0.003, -0.001] |
| First-session logical score (centred) | 0.026 | 0.008 | 3.35 | < 0.001 | [0.011, 0.041] |
| Spaced vs Massed (same model, Massed as reference) | 0.058 | 0.035 | 1.65 | = 0.098 | [-0.011, 0.127] |

**Confidence, correctly recognised trials only (LMM; no recognition term)**

| Term | b | SE | z | *P* | 95% CI |
|---|---|---|---|---|---|
| Intercept | 4.363 | 0.056 | 77.55 | < 0.001 | [4.253, 4.473] |
| Massed vs Once | 0.335 | 0.031 | 10.71 | < 0.001 | [0.274, 0.396] |
| Spaced vs Once | 0.418 | 0.032 | 13.18 | < 0.001 | [0.356, 0.481] |
| CET-4 (centred) | -0.001 | 0.002 | -0.31 | = 0.755 | [-0.004, 0.003] |
| Massed vs Once × CET-4 (centred) | 0.001 | 0.001 | 1.26 | = 0.209 | [-0.001, 0.003] |
| Spaced vs Once × CET-4 (centred) | 0.001 | 0.001 | 0.59 | = 0.553 | [-0.001, 0.002] |
| First-session study duration (centred) | -0.003 | 0.000 | -5.46 | < 0.001 | [-0.004, -0.002] |
| First-session logical score (centred) | 0.026 | 0.007 | 3.67 | < 0.001 | [0.012, 0.039] |
| Spaced vs Massed (same model, Massed as reference) | 0.083 | 0.030 | 2.77 | = 0.006 | [0.024, 0.142] |

*Note*: Step-1 condition-effect estimates were highly similar (final logical score S vs O b=2.53; recognition S vs O b=2.08); that is, condition effects remained similar after adjustment for first-session study duration and first-session logical score. The two confidence blocks are the recognition-centred model (overall adjusted contrast) and the model fitted to correctly recognised trials only (Supplementary Results S4; Supplementary Fig. 3). CET-4 moderation terms (Step 3 interactions) are reported in Supplementary Table 5.

#### Supplementary Table 5. CET-4 Moderation of the Spaced-versus-Massed Contrast in Step-3 Trial-Level Models

| Outcome (model) | S-vs-M × CET-4, *b* | *P* |
|---|---|---|
| Final logical score (LMM) | -0.00301 | 0.375 |
| Recognition (GEE, logit) | -0.01533 | <0.001 |
| Confidence (LMM) | -0.00046 | 0.665 |

*Note*: CET-4 scores were grand-mean centred. Each coefficient indexes the change in the spaced-versus-massed contrast associated with a one-point increase in CET-4; negative coefficients indicate that the spacing advantage decreased as CET-4 increased. Final logical score and confidence were analysed with LMMs including participant-specific random intercepts; recognition was analysed with a binomial-logit GEE and exchangeable within-participant correlation. All models adjusted for first-session study duration and first-session logical score; the confidence model additionally included recognition and the condition × recognition interaction. Values are direct Step-3 model coefficients and two-sided, unadjusted Wald *p* values; the moderation analyses were exploratory.

#### Supplementary Table 6. TFCE-corrected NPS inference over the bilateral central domain

| Family | Direction | Observed TFCE peak | Minimum corrected p | Corrected cells | Supported window centres (ms) | Supported sensors |
|---|---|---|---|---|---|---|
| path a | a < 0 | FT8, 590 ms | 0.0221 | 58 | 520–740 | FC4, FC6, FT8, C4, C6, T8 |
| path a | a > 0 | FC5, 380 ms | 0.9995 | 0 | none | none |
| path ab | NN: a < 0, b < 0, ab > 0 | T8, 660 ms | 0.0226 | 20 | 620–730 | C6, T8 |
| path ab | NP: a < 0, b > 0, ab < 0 | CP5, 350 ms | 0.1079 | 0 | none | none |

*Note.* The tested domain comprised 18 bilateral central searchlight centres (nine per hemisphere) and 71 window centres from 50 to 750 ms (100-ms width, 10-ms step), giving 1,278 sensor–time cells per direction. TFCE used H = 2 and E = 1, with within-hemisphere Delaunay adjacency, no cross-hemisphere spatial edges, and adjacency between successive windows. Path a and path ab were distinct correction families: a < 0 and a > 0 shared the path-a family; NN and NP shared the path-ab family. For each of 1,991 within-participant S/M label permutations with target blocks intact, each family's null value was the largest raw TFCE value across both hemispheres, all cells and its two directions. Corrected p = (1 + k)/1,992, where k counts null maxima at least as large as the observed TFCE value; support required $P \leq 0.05$. No correction was applied across families. Displayed p values are rounded to four decimal places after thresholding. Peak locations identify observed directional TFCE maxima, including for directions without corrected support; a listed peak does not itself establish significance. Reported spans are window-centre ranges, not continuous support at every listed sensor. All supported cells were right-central. Path-ab inference was not screened by path-a significance. Participant means of spaced-minus-massed NPS over the 20 NN cells averaged −0.007 r, positive in 11 of 28 participants (Figure 4e, bottom).

---

#### Supplementary Table 7. Corrected path-a cells by window centre

| Window centre (ms) | n | Supported sensors |
|---|---|---|
| 520 | 1 | C4 |
| 530 | 1 | C4 |
| 540 | 2 | FC4, C4 |
| 550 | 2 | FT8, C4 |
| 560 | 4 | FC4, FC6, FT8, C4 |
| 570 | 5 | FC4, FC6, FT8, C4, T8 |
| 580 | 6 | FC4, FC6, FT8, C4, C6, T8 |
| 590 | 5 | FC4, FC6, FT8, C6, T8 |
| 600 | 4 | FC4, FC6, FT8, T8 |
| 610 | 2 | FC6, FT8 |
| 620 | 2 | FC6, FT8 |
| 630 | 2 | FC6, FT8 |
| 640 | 2 | FC6, FT8 |
| 650 | 3 | FC6, FT8, T8 |
| 660 | 4 | FC6, FT8, C6, T8 |
| 670 | 4 | FC6, FT8, C6, T8 |
| 680 | 3 | FC6, C6, T8 |
| 690 | 1 | T8 |
| 700 | 1 | T8 |
| 710 | 1 | T8 |
| 720 | 1 | T8 |
| 730 | 1 | T8 |

| Window centre (ms) | n | Supported sensors |
|---|---|---|
| 740 | 1 | T8 |

*Note.* n is the number of searchlight centres with corrected a < 0 support at each listed window centre. Counts sum to 58 sensor–time cells, all right-central, within the tested domain of 18 bilateral central centres × 71 window centres (50–750 ms; 1,278 cells). Each estimate represents a 100-ms window indexed by its centre, with successive centres 10 ms apart. Only window centres with at least one supported cell are listed; unlisted tested window centres and all left-central centres had no corrected a < 0 support. Support follows the path-a family in Supplementary Table 6: 1,991 intact-target-block label permutations and one shared raw-maximum TFCE null across both hemispheres, all cells and both signs of a, with corrected $P \leq 0.05$. No a > 0 cell survived correction.

---

#### Supplementary Table 8. Corrected NN indirect-effect cells by window centre

| Window centre (ms) | n | Supported sensors |
|---|---|---|
| 620 | 1 | T8 |
| 630 | 1 | T8 |
| 640 | 2 | C6, T8 |
| 650 | 2 | C6, T8 |
| 660 | 2 | C6, T8 |
| 670 | 2 | C6, T8 |
| 680 | 1 | T8 |
| 690 | 1 | T8 |
| 700 | 2 | C6, T8 |
| 710 | 2 | C6, T8 |
| 720 | 2 | C6, T8 |
| 730 | 2 | C6, T8 |

*Note.* n is the number of searchlight centres with corrected NN support at each listed window centre. NN denotes a < 0, b < 0 and ab > 0. Counts sum to 20 sensor–time cells, all right-central: T8 at 620–730 ms (12 cells) and C6 at 640–670 and 700–730 ms (8 cells). The tested domain comprised 18 bilateral central centres × 71 window centres (50–750 ms; 1,278 cells per direction). Each estimate represents a 100-ms window indexed by its centre; successive centres are 10 ms apart. Only window centres with at least one supported NN cell are listed. Support follows the path-ab family in Supplementary Table 6: 1,991 within-participant S/M label permutations with target blocks intact and one shared raw-maximum TFCE null across both hemispheres, all cells and the NN and NP directions, with corrected $P \leq 0.05$. No NP cell survived correction on either side, and no left-central NN cell survived correction. Path-ab inference was corrected separately from path-a inference, without screening by path-a significance; 11 of the 20 NN-supported cells also had corrected a < 0 support.

#### Supplementary Table 9. Representative stimulus items

Each target word is decomposed into familiar cognate components (first association) and an etymological derivation linking those components to the meaning (second association), illustrating the two-stage meaningful-learning structure.

| Target word | Structural decomposition (first association) | Etymological derivation (second association) | Meaning |
|---|---|---|---|
| libretto | *liber* = book/paper (cf. *library*); *-etto* = diminutive suffix | a small collection of pages → a collection of song lyrics | n. libretto (song-lyric collection) |
| oration | *-orat-* = to speak (cf. *oral*); *-ion* = noun suffix | something spoken aloud → a speech | n. oration |
| apparel | *ap-* = *ad-* (toward); *par* = to prepare (cf. *prepare*) | to get ready → to prepare one's clothing | n. apparel |
| ballad | *ball* = to dance; *-ad* = noun suffix | a song played while dancing → a folk song | n. ballad |
| crusade | *crus-* = cross (cf. *cross*), from the cross carried by soldiers | the Crusades → a reform movement | n. crusade / reform movement |
| cyclone | *cycl-* = circle (cf. *cycle*) | wind spinning in a circle → a whirlwind | n. cyclone |

#### Supplementary Table 10. Window-centre ranges with corrected positive CORnet-Z RSA support

| CORnet-Z stage | Spaced (S) | Once-learning (O) | Massed (M) |
|---|---|---|---|
| V1 | — | 40–230 | 170–210 |
| V2 | — | — | 170–210 |
| V4 | — | 150–250 | — |
| IT | 420–460 | 90–250 | 330–510 |

*Note.* Values are window centres in milliseconds. Each interval contains consecutive window centres at which at least one of the 17 posterior searchlight centres showed positive RSA at TFCE-corrected $P < 0.05$ (N = 28; 5,000 whole-participant sign flips). Family-wise error was controlled separately within each condition × stage's 17-centre ×

91-window field, not across the 12 families. Windows were 100 ms wide and advanced in 10-ms steps. A dash indicates no TFCE-corrected positive RSA support within that family, not evidence that the representation is absent. Different centres may contribute to different portions of an interval; the intervals do not indicate support at every centre, for the spatially averaged curve, or in every participant. Their boundaries and lengths are not estimates of processing onset, offset or duration. Direct between-schedule contrasts yielded no corrected cell (Supplementary Methods SM12; Supplementary Results S6).
